# Circuit mechanisms to transform neural population dynamics for motor control

**DOI:** 10.1101/2025.02.21.639459

**Authors:** David J. Herzfeld, Stephen G. Lisberger

**Affiliations:** Department of Neurobiology, Duke University School of Medicine, Durham, NC, 27710, USA; Department of Neuroscience, University of Wisconsin-Madison, Madison, WI, 53705, USA

**Keywords:** neuron type, smooth pursuit, learning, basis set, temporal decomposition, mossy fiber, unipolar brush cell, Golgi cell, granule cell, circuit computation

## Abstract

We identify the input-output transformations performed by the cerebellar circuit and isolate the circuit properties that implement them. Extracellular recordings exploited new technology for neuron-type identification in part of the cerebellar cortex that is causal for an oculomotor behavior whose neural substrate is well-understood in monkeys. Data demonstrate how multiple classes of interneurons in the cerebellar circuit cooperate to transform incoming mossy fiber responses into Purkinje cell outputs. Computer models suggest that temporal decomposition in the granule cell layer enables flexible reconstruction of transformed output dynamics, regulated by cerebellar plasticity, to convert eye position-like inputs into predominantly velocity-driven Purkinje cell outputs. Convergence of mossy fiber inputs with diverse direction preferences would increase input layer dimensionality and enable learning of highly flexible direction tuning. The neural circuit mechanisms suggested by our data outline a cerebellar operating principle that enables flexible input-output transformations and could generalize across cerebellar and other brain areas.

## Introduction

Understanding how neural circuits process information to generate behavior is a fundamental goal of systems neuroscience. Circuit computations can be envisioned as multiple transformations^1^ of neural population dynamics^2,3^ - from sensory inputs^4^ to intermediate representations^5^ to the ultimate drive of precise muscle activations^6^. The cerebellum provides an ideal platform for understanding circuit mechanisms that transform population dynamics. The cytoarchitecture of the cerebellum is highly conserved^7,8^, with a well-characterized circuit organization^9^, discrete neuron types^9,10^, and crucially, well-defined inputs and a single neural population output^11,12^. It has long been thought that the cerebellar circuit performs a canonical, repeated computation^13,14^ across its different regions for both motor and non-motor behaviors. Thus, identifying the circuit mechanisms that transform population dynamics in one region of the cerebellum may generalize across cerebellar regions and tasks.

We focus on transformations of population dynamics in the cerebellar floccular complex, a structure chosen for analysis because of the unique interpretational power provided by extensive knowledge about its neural signal processing^15^ and causal connection to a quantifiable behavior^16,17^. The floccular complex is essential^17,18^ for controlling smooth pursuit eye movements via disynaptic connections to extraocular motor neurons^19^. Prior recordings from its Purkinje cell outputs^20–22^ and mossy fiber inputs^23,24^ tentatively characterized the input-output transformation of the floccular circuit. Now our goal was to understand how the complete circuit performs those input-output transformations. The circuit is known^9^: incoming mossy fibers synapse onto granule cells, whose parallel fiber axons provide excitatory input to Purkinje cells, the sole output of the cerebellar cortex. Additional neuron types^10^, including Golgi cells, unipolar brush cells, and molecular layer interneurons (among others^25,26^), use both feed-forward and recurrent connections^11^ to modulate the activity within the pathway between mossy fibers and Purkinje cells.

We take advantage of recent advances in large-scale multi-contact electrode technology^27,28^ and neuron-type identification strategies^29,30^ to record populations of identified cerebellar input fibers, output neurons, and interneurons during motor behavior. We reveal neural circuit mechanisms that can transform cerebellar inputs into outputs in both time and space through two specific computational motifs: temporal decomposition in the granule cell layer and spatial reorganization in the granule cell and molecular layer. We show that the same computational motifs can be generalized for a wide variety of transformations, potentially throughout the cerebellum and in many neural circuits across the brain.

## Results

### Input-output transformations within the floccular complex during smooth pursuit eye movements

We first document directional and temporal input-output transformations in the floccular complex and reveal that they are partitioned along neuron-type boundaries. Based on the data, we suggest that the temporal transformation occurs in the granule cell layer and the directional transformation occurs through the interaction of parallel fiber and molecular layer interneuron (MLI) inputs to Purkinje cells in the molecular layer. Computational analyses instantiate the proposed details of both transformations.

Using multi-contact probes, we recorded extracellular action potentials from the floccular circuit (Fig. 1a) while monkeys performed smooth pursuit eye movements (Fig. 1b). We successfully identified all major neuron types schematized in Fig. 1a, with the exception of granule cells, using expert criteria^30^ (see ***Methods***), validated via ground-truth optogenetic identification in mice^29^.

**Fig. 1.**
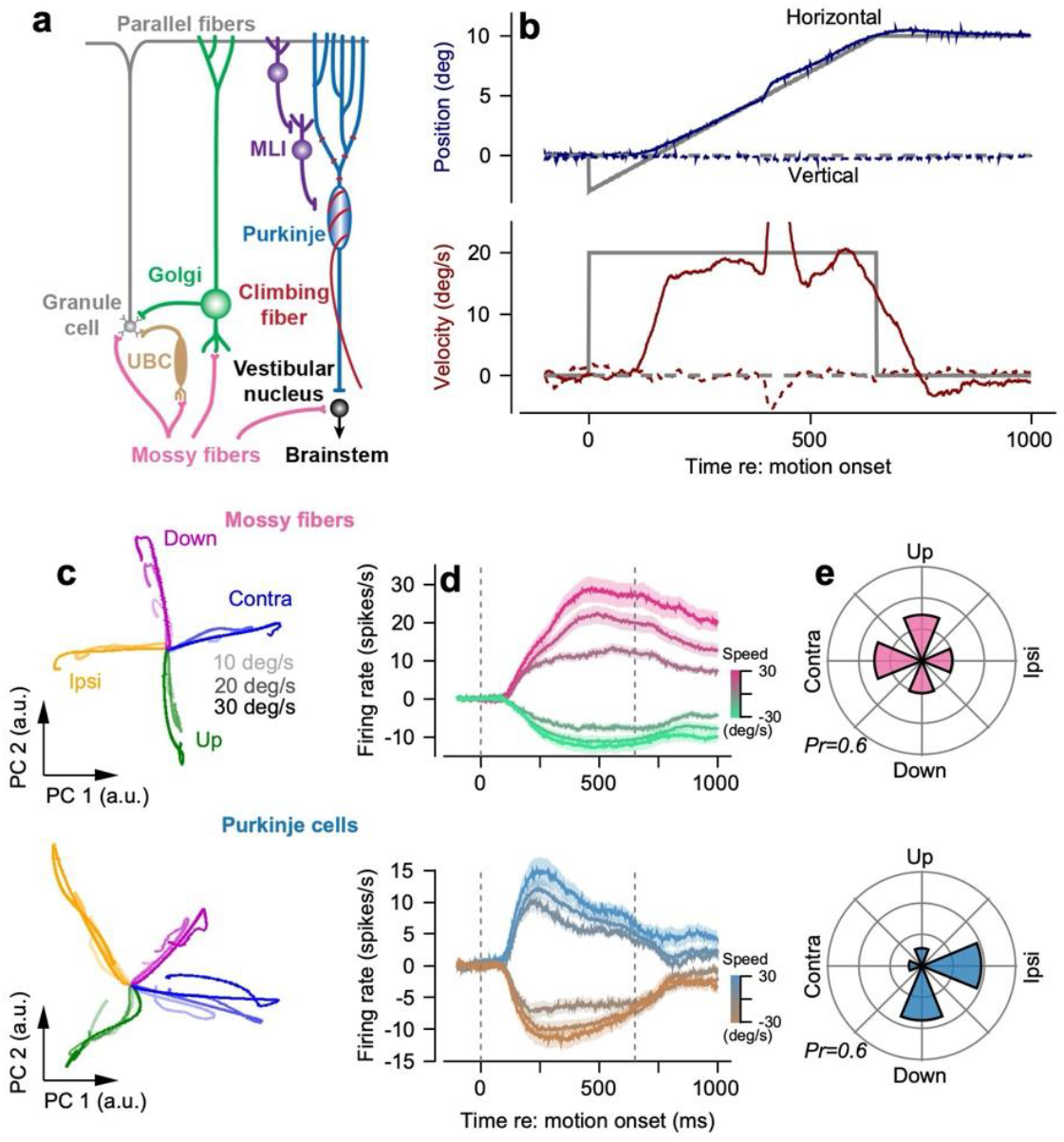
Temporal and directional input-output transformations performed by the cerebellar floccular complex during smooth pursuit. **a**, Simplified schematic of the cerebellar circuit in the floccular complex. **b**, Eye and target position (top) and velocity (bottom) versus time for an example smooth pursuit trial where the target moved exclusively in the horizontal direction. Monkeys fixated a central dot for a randomized interval (400-800 ms) before the target started to move at a constant velocity for 650 ms. Monkeys tracked the target throughout its motion and maintained eccentric fixation for 350 ms to receive fluid reward. Targets moved at 10, 20, or 30 deg/s along the four cardinal directions. **c**, Principal components computed separately for mossy fibers (top) and Purkinje cells (bottom) across target speeds (10, 20, 30 deg/s, increasingly saturated traces) and cardinal directions. **d**, Mean population responses of mossy fibers and Purkinje cells in their preferred and anti-preferred directions, relative to their respective baseline responses. Shaded bands represent mean ± SEM across neurons. **e**, Polar plots showing the distribution of preferred directions across the populations of mossy fibers and Purkinje cells. Preferred directions are defined as pursuit direction with maximal mean positive deviation of firing rate (0 to 650 ms after target motion onset) relative to the pre-trial baseline response. We note that the difference in direction tuning of Purkinje cells between **c** and **e** is due to the sign-agnostic nature of principal component analysis along with the comparable magnitudes of Purkinje cell modulation in the preferred and anti-preferred directions.

The inputs and outputs from the floccular complex, mossy fibers and Purkinje cells, follow distinct low-dimensional neural state-space trajectories^2,31^ (Fig. 1c). Mossy fiber trajectories align closely with the horizontal and vertical axes of the eye and terminate eccentrically in state-space with magnitudes that scale with pursuit speed. In contrast, Purkinje cell trajectories show principal axes that are rotated relative to the cardinal axes and return near the origin at movement completion. For both neuron types, the first two dimensions captured a substantial portion of the response variance: 38% and 29% for mossy fibers and 23% and 20% for Purkinje cells (a considerable amount given that we are accounting for responses in 4 directions by 3 speeds of target motion). The first two components dominate the responses: for mossy fibers and Purkinje cells, respectively, the 3^rd^ principal component accounts for only 7% and 11% of the variance.

The mean population responses of mossy fibers and Purkinje cells in the preferred and opposite directions reveal temporal dynamics that explain the difference in state-space trajectories (Fig. 1d). Mossy fiber responses resembled eye position traces, with persistent firing at movement termination that scaled approximately linearly with pursuit speed (preferred direction R^2^ = 0.83 ± 0.03). Purkinje cell responses peaked approximately 200 ms after motion onset with magnitudes that increased with pursuit speed and returned to baseline at the end of pursuit irrespective of final eccentric eye position. The shift from sustained mossy fiber responses to transient Purkinje cell responses requires the cerebellar circuit to perform a “temporal transformation.”

Directional tuning properties also differed between mossy fibers and Purkinje cells (Fig. 1e). Mossy fibers showed uniformly distributed preferred directions across the cardinal axes (circular dispersion: *<u>R</u>*= 0.13) with essentially no modulation of firing for pursuit orthogonal to their preferred axis. Purkinje cells’ preferred directions were more strongly biased, with preferred directions close to ipsiversive and downwards (*<u>R</u>*= 0.58)^21,22,24,32–34^, requiring the cerebellar circuit to perform a “directional transformation.” Purkinje cells exhibited diverse directional tuning, with varying degrees of inhibition in the anti-preferred direction and often small increases or decreases in firing in orthogonal directions.

Different neuron types exhibit distinct temporal response properties, suggesting each plays a unique computational role in shaping cerebellar signals (Fig. 2a). For pursuit in their preferred directions, the responses of mossy fibers and unipolar brush cells were related to both eye position and velocity and remained elevated during eccentric fixation at pursuit termination.

Molecular layer interneurons were related to eye velocity and Purkinje cells to eye velocity and acceleration, with firing rates that returned to near baseline at pursuit termination while the monkey fixated eccentrically. Golgi cells showed minimal modulation during pursuit.

**Fig. 2.**
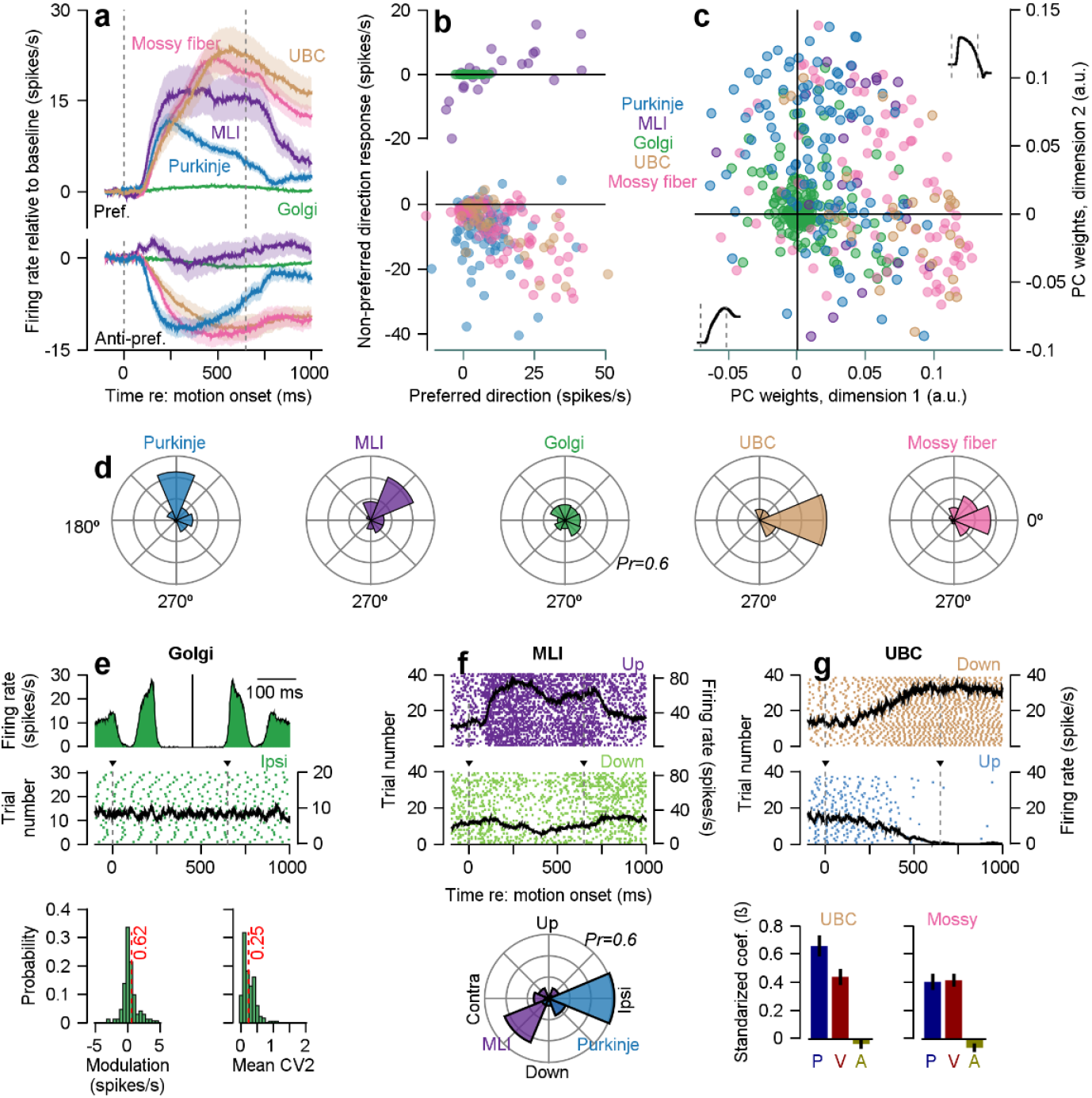
Functional organization and response heterogeneity across cerebellar neuron types during smooth pursuit eye movements. **a**, Mean modulation of firing rate as a function of time for cerebellar neurons, segregated by neuron type, during 20 deg/s pursuit in each neuron’s preferred (top) and anti-preferred (bottom) pursuit directions. Error bands show mean ± SEM across neurons. **b**, Scatter plot of the relationship between responses in the preferred and non-preferred direction. Each dot corresponds to an individual identified neuron, colored by neuron type. **c**, Scatter plot of the loadings onto the first two principal components computed across all identified floccular neurons. Principal components were computed across all identified neurons using only their preferred direction responses. Permutation analysis revealed that all neuron type centroids were significantly different from a neuron-type agnostic distribution (mossy fibers, *n = 86*, p = 10^−4^; unipolar brush cells (UBCs), *n = 30*, p = 10^−4^ Golgi cells, *n = 186*, p = 10^−4^; Purkinje cells, *n = 101*, p = 10^−4^; molecular layer interneurons (MLIs), *n = 23*, p = 0.039). The inset traces at the end of each axis show the two principal components. **d**, Distribution of angular locations, irrespective of magnitude, for each neuron type derived from the scatter plot shown in **b. e**, Golgi cell features. From top to bottom: autocorrelogram of an exemplar Golgi cell; raster plot of the same Golgi cell aligned to the onset of ipsiversive target motion where the black curve shows the mean firing rate across all trials shown in the raster plot; probability distributions showing the modulation in the preferred pursuit direction and the CV2 of all Golgi cells computed across complete recording sessions. Red dotted lines denote population means across Golgi cells. **f**, MLI features. From top to bottom: raster plots for an exemplar MLI in its preferred and anti-preferred pursuit directions, where black curves denote the mean firing rate of the MLI across the trials shown in the raster; polar probability distributions of preferred directions for molecular layer interneurons (purple) and simultaneously-recorded Purkinje cells (blue). The mean modulation of the molecular layer interneuron population was 12.2 ± 3.0 spikes/s in their preferred direction, but 0.4 ± 1.6 spikes/s in their anti-preferred direction. The preferred direction of MLIs differed from that of their connected Purkinje cells by −121.8° ± 10.4° (circular mean ± SEM). **g**, UBC features. From top to bottom: rasters for an exemplar UBC in its preferred (downwards) and anti-preferred (upwards) pursuit directions, where black curves denote mean firing rates during pursuit across the trials plotted in the raster; bar graphs to compare standardized coefficients relating eye position (P), velocity (V) and acceleration (A) to firing rates of UBCs (left) and mossy fibers (right) for pursuit in each neuron’s preferred direction at 20 deg/s. Shaded regions and error bars in all panels denote mean ± SEM across neurons. Extended Data Fig. 1 documents the responses of our full samples of Golgi cells, MLIs, and UBCs in their preferred directions.

The features of preferred versus non-preferred mean responses (Fig. 2a) were consistent across individual neurons of each type (Fig. 2b). Purkinje cells, mossy fibers, and unipolar brush cells showed reciprocal responses (negative values in the anti-preferred direction). Golgi cells showed minimal modulation in either preferred or anti-preferred directions, while most MLIs exhibited positive, non-reciprocal modulation in both directions.

Principal component analysis of trial-averaged pursuit responses in each neuron’s preferred direction at 20 deg/s showed progressive changes in temporal responses across the cerebellar circuit. The two dominant components together explained 81% of the population variance (49% and 32%, respectively). The first principal component (Fig. 2c, x-axis inset) resembled eye position, with a near monotonic increase over time and sustained deviation. The second principal component (Fig. 2c, y-axis inset) exhibited an early peak followed by a return to near baseline, resembling a combination of eye velocity and acceleration. The third principal component accounted for 2.5% of the variance.

In the two-dimensional space defined by the first two principal components (Fig. 2c), Golgi cells are concentrated near the origin. Purkinje cells show near zero weights in the first dimension but consistently positive weights in the second dimension. Mossy fibers and unipolar brush cells show the opposite trend, with weights centered near zero in the second dimension but strongly positive in the first dimension. Much of the within-neuron-type variation in the spatial distribution shown in Fig. 2c results from heterogeneity in response magnitude rather than differences in temporal properties within neuron types. Angular distributions in the two-dimensional space (Fig. 2d) differ across neuron types, reflecting distinct relative contributions of the two principal components to neuron firing.

Before turning to computational analysis, we draw conclusions based on the data in Fig. 2. Specifically, features of each cerebellar interneuron type suggest that they could not contribute to the temporal input-output transformation within the cerebellar circuit, as well as how they might contribute to the directional transformation.

- *Golgi cells* showed essentially no temporal modulation of firing during ipsiversive pursuit with regular firing at low rates (mean rate = 12 spikes/s, mean “CV2^35^” = 0.25 ± 0.01) (Fig. 2e). We conclude that Golgi cells provide tonic but temporally and directionally (Fig. 2b, c) unmodulated inhibition of granule cells.
- *Molecular layer interneurons* showed preferred directions opposite to those of their connected Purkinje cells. Thus, a molecular layer interneuron’s firing, related primarily to eye velocity in its preferred direction (Fig. 2f), provides inhibition that suppresses the firing of Purkinje cells in their anti-preferred direction. The presence of velocity signals in both molecular layer interneurons and Purkinje cells argues for a shared upstream transformation available to both populations in parallel, rather than independent intrinsic mechanisms reimplemented in each cell type. Molecular layer interneurons displayed minimal modulation of firing rate during pursuit in their anti-preferred direction (Fig. 2b) and therefore have little impact on the temporal dynamics of Purkinje cell preferred-direction responses but possibly greater impact on their direction tuning.
- *Unipolar brush cells* showed clear positive and negative modulation in their preferred and anti-preferred directions (Fig. 2b), had fully sustained modulation in both directions after pursuit termination and, like mossy fibers, responded primarily to a combination of eye position and velocity (Fig. 2g). Regression analysis showed stronger eye position encoding for unipolar brush cells compared to mossy fibers (histograms in Fig. 2g, standardized coefficient, β = 0.65 ± 0.07 versus 0.40 ± 0.06, t(114) = −2.36, p = 0.02). The dominant eye position response of unipolar brush cells suggests that they are not part of the transformation to velocity-related Purkinje cell or molecular layer interneuron responses.

Finally, the temporal transformation cannot be explained by synchrony in multiple mossy fibers inputs to granule cells because we found no evidence for mossy fiber synchrony other than the amount expected from covariation of mossy fiber firing rates during pursuit (Extended Data Fig. 2).

### A data-driven computational approach for predicting granule cell temporal representations

The responses across neuron types imply that the temporal input-output transformation in the floccular complex occurs in the granule cell layer. Given the inaccessibility of granule cells to extracellular recordings^29^, we adopted a computational approach: we trained a deep-learning model of the mossy fiber-to-Purkinje cell pathway with explicit granule cell dynamics (Fig. 3a). Our goal was to discover the crucial computational features necessary for successful temporal transformation before mapping the successful computational features onto potential cerebellar circuit and cellular mechanisms.

**Fig. 3.**
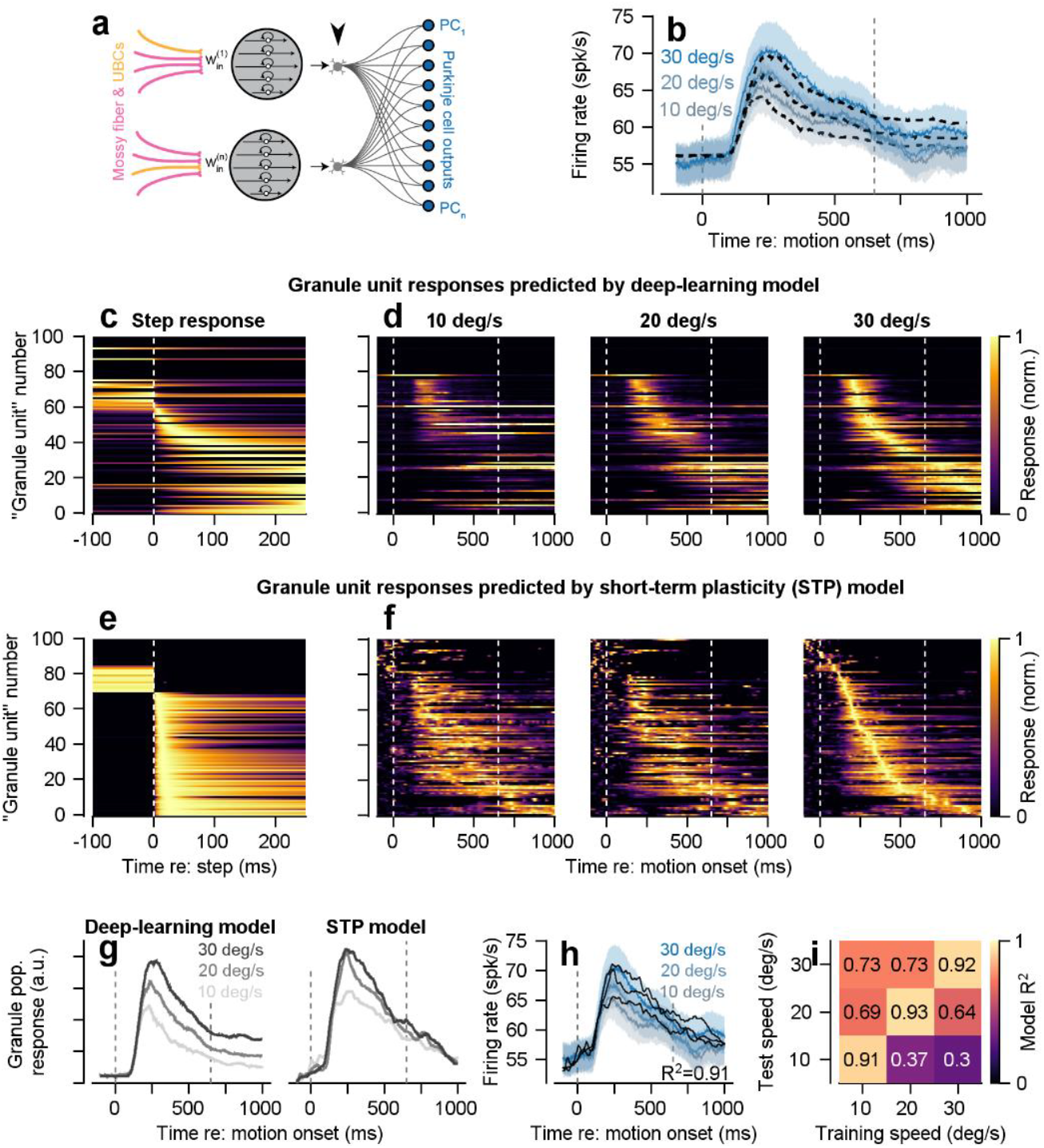
Models reveal a potential mechanism of the temporal transformation between mossy fibers and Purkinje cells. **a**, Deep-learning model that transforms mossy fiber and unipolar brush cell inputs to create “granule unit” (black arrowhead) responses that can be combined with linear weights to reproduce Purkinje cell firing rates. **b**, Success of the model shown in **a** to account for Purkinje cell data. Dashed black traces show mean model responses to three different speeds of target motion. Blue traces show population mean Purkinje cell firing rates across speeds. Note that models were fitted to individual Purkinje cell responses and the results averaged for visualization. **c**, Responses of granule units in the deep-learning model to a step change in the input at *t=0* ms. Each unit’s response was normalized to its maximum firing rate. Rows are ordered by the time of the peak response. **d**, Responses of granule units in the deep-learning model to the three pursuit speeds shown in **b**. Each granule unit’s response was normalized to its peak response across all three pursuit speeds and ordered based on the timing of the peak response to 30 deg/s target motion (far right). **e**, Responses of granule units to a step input of mossy fiber responses in a model based on short-term plasticity (STP) at the mossy fiber to granule unit synapses. **f**, Responses of granule units in the STP model to four randomly sampled mossy fiber inputs across pursuit speeds. **g**, Population responses of granule units in the deep-learning and STP models. **h**, Fits (black traces) to the mean responses of Purkinje cells across pursuit speeds (colored traces) using the LTP/LTD rule described in the Methods to adjust the weights of granule units from the STP model on Purkinje cells. **i**, Generalization matrix showing STP model R^2^ when trained on the data for one target speed at a time and tested on all 3 target speeds.

Inputs to each granule unit comprised weighted single-trial responses from 4 randomly selected mossy fibers or unipolar brush cells. Here, each granule unit comprises 25 single time constant filters, with shared time constants, but unique filter weights across granule units. Thus, each granule unit has a “personalized” temporal response to its inputs. Granule unit output weights were trained to predict the time-varying firing rates of each recorded Purkinje cell. The resulting granule unit basis set had dynamics that allowed the model Purkinje cells to reproduce the mean temporal profiles of recorded Purkinje cell responses across a range of pursuit speeds (Fig. 3b, R^2^ = 0.93). Note that here, and throughout the paper we fit models to each individual Purkinje cell in our sample and report the mean time-varying responses and the average R^2^ across Purkinje cells and pursuit speeds (not the R^2^ of the average responses).

The model network temporally decomposed its mossy fiber and unipolar brush cell inputs to create a range of transient responses where the degree of filtering and subsequent duration of granule unit firing were temporally linked. When the model was supplied with a step input, some units produced transient, high-pass filter like responses, while others showed sustained, low-pass filter like activity (Fig. 3c). The latency of each responsive granule unit’s peak response was correlated to its response duration (correlation test between rise and decay time constants, t(36) = 6.6, p < 10^−6^). The basis set showed similar patterns of units that we could characterize as high-pass and low-pass when we drove the model with mean rather than single-trial inputs (Fig. 3d).

### A biologically motivated model of granule cell temporal transformations

We used the evidence for short-term plasticity (STP) in the mossy fiber to granule cell synapses^36,37^ as the basis for a model that could create a temporally-decomposed granule unit basis set using biological principles. To keep the model computationally tractable, we extended the classic Tsodyks-Markram STP framework^38^ to include a dual transmitter pool. Each granule unit receives inputs from 4 randomly-weighted mossy fiber/unipolar brush cells and implements STP through a “Dual Pool Synapse” with (1) a “fast” pool of neurotransmitter that is readily releasable and (2) a “slow” pool of neurotransmitter that replenishes the fast pool (see ***Methods***).

The resulting granule units exhibited a broad range of temporal filtering properties that agreed well with previously proposed “spectral timing” models of granule cell activity^39–41^. The step-responses of model granule units also agreed with those predicted by the deep-learning model (Fig. 3e), as did the model’s responses to mossy fiber and unipolar brush cell inputs during pursuit (Fig. 3f). The STP model showed emergent correlations between rise and decay times (t(49) = 2.2, p = 0.03), consistent with the deep-learning model. Other emergent properties of the STP granule unit basis set are (1) granule unit population activity (Fig. 3g) increased with pursuit speed to allow Purkinje cell modulation to increase with speed and (2) granule unit timing was consistent across pursuit speed to enable generalization across pursuit speeds.

The mean response of a population of 100 granule units (Fig. 3g) resembled the mean of the temporal profiles observed in experimentally recorded Purkinje cells (Fig. 3h, blue curves). Thus, it was not surprising that optimizing the output weights of the granule units via the LTP/LTD learning rule provided an excellent fit to the responses of our population of Purkinje cells (Fig. 3h, mean R^2^=0.91). Further, the STP basis set generalized well when trained on the responses to a single target speed and tested on the other 2 target speeds (Fig. 3i), Again, note that we fitted each Purkinje cell individually and we report averages of the time varying firing rates and the mean R^2^ for simplicity.

Analysis of other plausible basis sets emphasized our main conclusions about the properties needed in the granule cell basis set: temporal decomposition of sustained inputs with scaling and consistent timing of each unit’s responses across target speeds. The other models share some of these features in a minority of their units and therefore are able to reproduce the time varying Purkinje cell firing to some degree, albeit less well than the STP basis set when using the LTP/LTP learning rule (Extended Data Fig. 3, 4). The other basis sets also perform reasonably, though less well than the STP basis set, when we used least squares optimization as the learning rule (Extended Data Fig. 5-7).

The details of our optimization procedure appear in the ***Methods***. In brief, for each granule unit model, we optimized the hyperparameters that bound granule unit parameters rather than the parameters themselves. For each optimizer iteration, we: (1) sampled random parameter values within these bounds, (2) computed each granule unit’s response to its 4 mossy fiber/unipolar brush cell inputs, (3) optimized parallel fiber-to-Purkinje cell weights to match recorded Purkinje cell firing, (4) calculated mean squared error, and (5) adjusted hyperparameters to potentially improve the next fit. Directly optimizing individual granule unit parameters would have yielded perfect fits with limited mechanistic insight.

### A cerebellar circuit model that reproduces the directional and temporal response properties of individual Purkinje cells and molecular layer interneurons

We subscribe to Feynman’s view: “What I cannot create, I do not understand”. Thus, our next step was to create a circuit model of the cerebellar cortex with realistic architecture and elements that show biomimetic neural responses.

Our model used weighted combinations of modeled granule units and measured responses of molecular layer interneurons to transform the temporal firing rate responses and direction tuning of mossy fibers and unipolar brush cells into the firing of individual Purkinje cells during pursuit eye movements. The model (Fig. 4a) consisted of 86 mossy fibers and 31 unipolar brush cells with the firing profiles from our recorded population, 1,000 granule units based on the STP model, 250 molecular layer interneurons (augmented from our recorded *n=23* population), and our 101 recorded Purkinje cells. Each granule unit used the STP model to transform input from 4 randomly-chosen mossy fibers and unipolar brush cells (Fig. 4a) into velocity-dominant, temporally decomposed, and directionally diverse granule unit activity (Fig. 4b). The convergence of randomly weighted inputs with different preferred directions created granule units with diverse and irregular direction tuning. The LTP/LTD learning rule (see ***Methods***) delivered excellent fits of both direction tuning and temporal dynamics of the 250 molecular layer interneurons (exemplar shown in the top row of Fig. 4c); the mean R^2^ for molecular layer interneurons was 0.71 (Fig. 4e). Note that the preferred directions of molecular layer interneurons were rotated by 180° to account for the opposite preferred directions relative to their Purkinje cell neighbors (Fig. 2f).

**Fig. 4.**
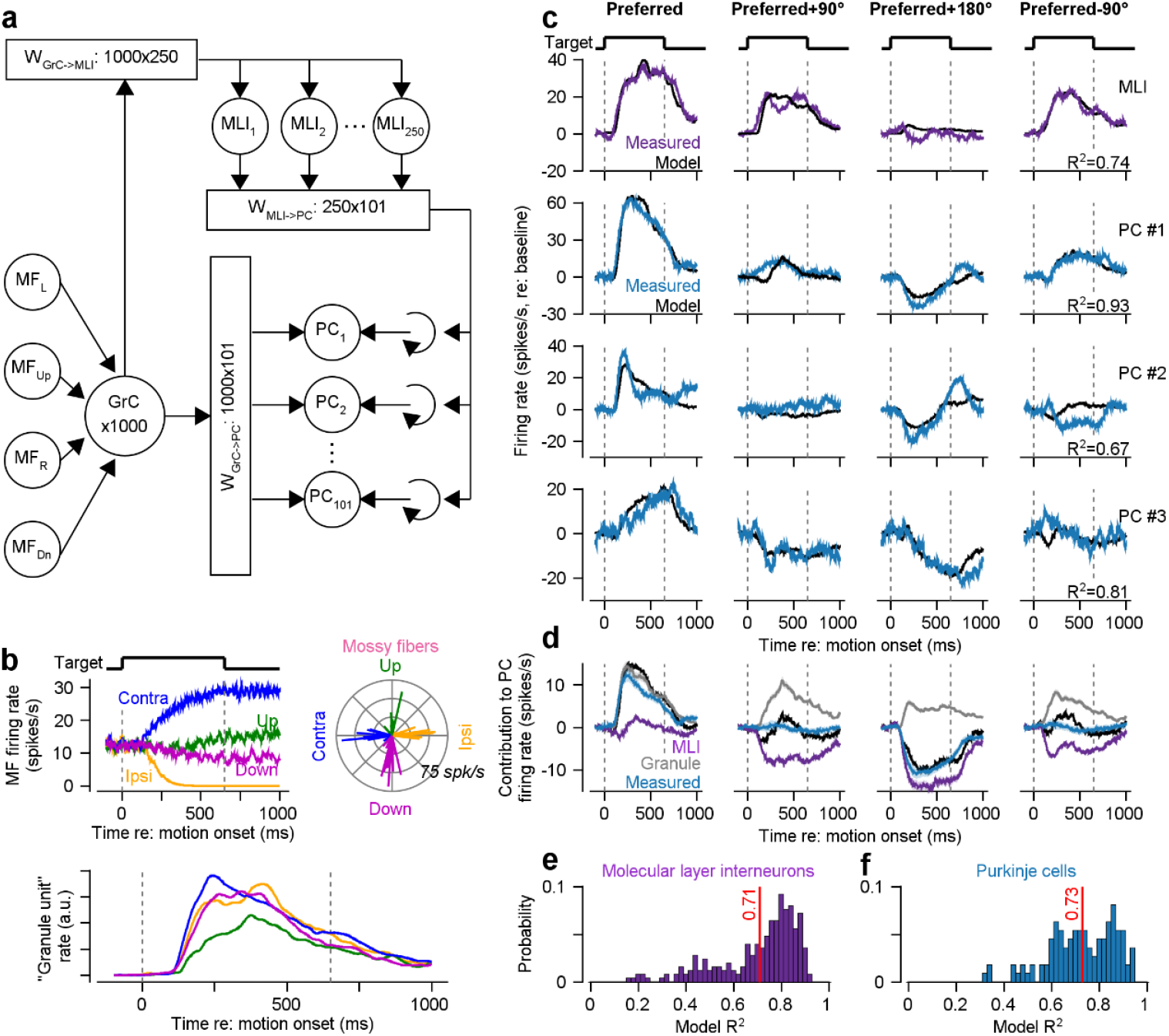
Model granule cell populations provide a sufficient basis for simulation of molecular layer interneuron and Purkinje cell responses. **a**, Schematic diagram of a circuit model of the cerebellar cortex used to predict the responses of individual molecular layer interneurons and Purkinje cells. **b**, Top left plot shows the responses of an example mossy fiber to 20 deg/s pursuit in the four cardinal directions. Right polar plot shows the response magnitude and preferred directions of mossy fibers Bottom row shows the responses of modeled STP granule units in the four cardinal directions. **c**, Fitted responses of an exemplar molecular layer interneuron (MLI, top row) and three exemplar Purkinje cells (rows 2-4) to 20 deg/s pursuit in four directions relative to each neuron’s preferred direction of pursuit. Black traces show model fits in all panels. **d**, Blue traces show mean modulation of Purkinje cell firing, black graces show mean model fits, and purple and gray graces show mean inputs from MLIs and granule units. **e-f**, Mean model R^2^s for molecular layer interneurons **(e)** and Purkinje cells **(f)**. Vertical red lines in **e-f** denote the means across the respective populations.

Proper weighting of the STP model of granule cell dynamics and the biomimetic molecular layer interneurons by the LTP/LTD learning rule accounted well for both the temporal dynamics and the direction tuning of the simple-spike activity of all individual Purkinje cells. Across the population of 101 Purkinje cells, the mean R^2^ was 0.73 (Fig. 4f). We could achieve almost perfect fits with a more-cleanly contrived granule unit basis set and least squares determination of weights, but we opted for more biologically relevant mechanisms.

We understood how the model works, and potentially how the cerebellar circuit computes, by analyzing the weighted contribution of granule cells and molecular layer interneurons to Purkinje cell firing across directions (Fig. 4d). In the preferred direction, Purkinje cell activity was driven almost entirely by excitatory input from the granule units. In the anti-preferred direction, the molecular layer interneuron population provided significant inhibitory input to counteract granule unit excitation. Along orthogonal axes, inputs from granule units and molecular layer interneurons approximately cancelled, resulting in Purkinje cell responses that were weakly modulated from baseline levels. The increased dimensionality of the representations of time and direction in the granule unit population allowed great flexibility in reproducing Purkinje cell responses.

### Emergent properties of the cerebellar circuit model

Any good model of the cerebellar circuit must (1) have emergent properties that account for previously described behavioral and neurophysiological observations and (2) generalize to other types of inputs found in other cerebellar regions.

The model in Fig. 4 reproduces at least two behaviors that were not built in: previous results documenting generalization of single-trial learning^42^ and the impact of the timing of the instructive stimulus^43^. Both cases depend on properties of the granule unit basis set in relation to time and eye velocity.

- To study generalization in monkeys, we induced learning with pursuit target motion at 20 deg/s and an instruction that started 250 ms later and provided orthogonal target motion at 30 deg/s for 450 ms^42^ (Fig. 5a). The instruction evokes complex-spike responses in the 100 milliseconds following the instruction (Fig. 5d, shaded region) and causes a well-timed depression of simple-spike firing rate (Fig. 5g). We measured learning in the subsequent test trial with slower or faster pursuit target motion (Fig. 5b). In our behavioral data^42^, eye movement deviation in the instruction direction of the test trial scaled with test target speed in the pursuit direction (Fig. 5e). In our model, we simulated the same experiment by reducing the connection weights from the subset of parallel fibers that were active at the time of instruction during pursuit at 20 deg/s (250 ms, red arrow) and measuring the learning-induced changes in Purkinje cell simple-spike output on the subsequent test trial for pursuit speeds of 10, 20, and 30 deg/s. The model predicted a learned depression in the Purkinje cell population response that scaled with pursuit target speeds (Fig. 5h), in agreement with our behavioral results. The scaling of firing rates with pursuit speed arises because the magnitude of granule cell population activity increases with pursuit speeds and a fixed amount of synaptic depression causes a post-synaptic response that scales with the change in firing of the pre-synaptic fibers.
- To study learned timing in monkeys, we repeated several hundred learning trials with a fixed interval of 150, 250, or 500 ms between the onset of target motion and the time of the instruction^43^ (Fig. 5c). Long intervals yielded smaller and temporally-broader behavioral learned responses (Fig. 5f). We simulated learned timing in our model by reducing the connection weights to Purkinje cells from parallel fibers that were active at one of the three intervals post-target motion onset: 150 ms, 250 ms, and 500 ms (colored arrows in Fig. 5f, i). The resulting changes in modeled Purkinje cell simple-spike responses (Fig. 5i) were consistent with the timing-dependent characteristics observed in learned eye movement responses (Fig. 5f). The dependence of modeled learned response on the timing of the instructive stimulus is due to positive correlation between the rise and decay time constants for granule layer units. Granule cells whose synapses are subject to plasticity for an early complex spike have temporally briefer responses than do the granule cells active later in the pursuit trial. Comparable temporal basis sets provide a possible neural mechanism for the learned timing in eye blink conditioning^44^.

**Fig. 5.**
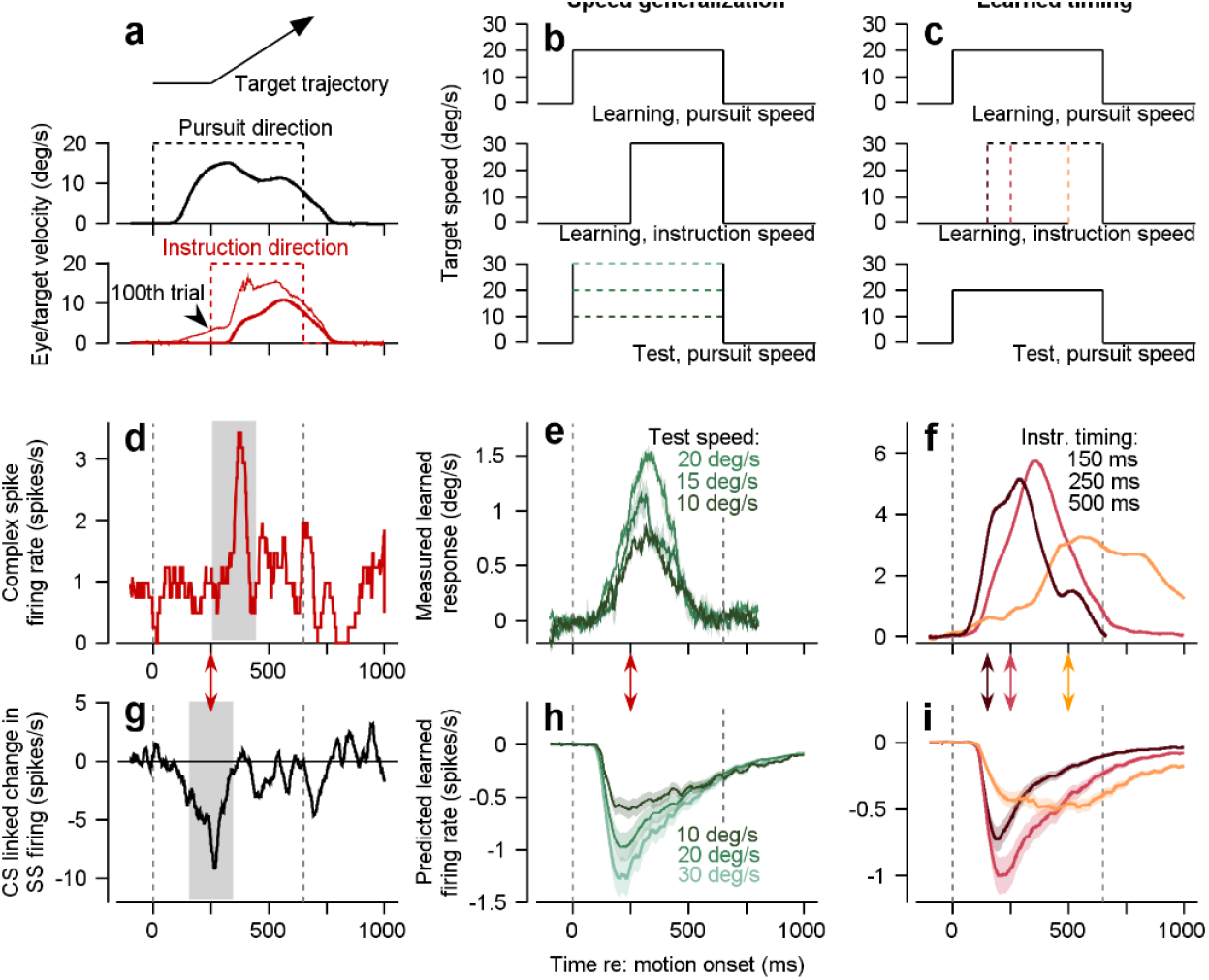
Emergent properties of the cerebellar circuit model during pursuit learning. **a**, Target position trajectory during a direction-learning trial (top). Eye and target velocity timeseries in the original pursuit direction (middle) and orthogonal instructive direction (bottom). Arrowhead in the bottom plot highlights learned eye velocity after 100 learning trials in the same direction. **b**, Target velocity profiles for a learning paradigm that tests speed generalization. Learning is instantiated as in **a** (top and middle traces). Speed generalization is probed in test trials (bottom) where target speed in the original pursuit direction is varied. **c**, Target velocity profiles for a pursuit learning paradigm that tests learned timing of the instructive stimulus. An instructive signal occurs either 150 ms, 250 ms, or 500 ms after target motion onset (middle). Learning is probed after 100 learning trials in a test trial with target motion only in the pursuit direction (bottom). **d**, Complex spike response measured during direction learning in trials where the instruction was in the preferred complex-spike direction (CS-on) of the Purkinje cell under study. Grey shaded region denotes instruction-linked complex-spike period. **e**, Eye velocity responses after learning, tested with probes of various speeds from **b**. Data adapted from reference ^42^. **f**, Eye velocity responses after 100 learning trials using different instruction timings, as in **c**. Data replotted from reference ^43^. **g**, Complex-spike-linked trial-over-trial change in simple-spike firing for the Purkinje cell shown in **d**. Grey shaded region highlights the well-timed simple-spike depression due to the occurrence of a complex spike following the instruction. **h**, Change in simple-spike responses predicted by the model for the paradigm of **b. i**, Change in simple-spike firing predicted by the model following learning trials with each of the three instruction timings from the experimental paradigm of **c**. Shaded regions in all panels denote mean ± SEM across neurons or behavioral learning paradigm replicates.

Other granule unit basis sets perform about as well as the STP basis set on speed generalization of learning, but quite poorly on learned timing (Extended Data Fig. 4 and 7).

### Generalization of the temporal transformation to other input-output modalities

The ability of a specific neural mechanism to account for the input-output transformation in one part of the cerebellum raises the question to whether it could have wider applicability. If so, it becomes plausible to think that a generalizable neural mechanism might work in other parts of the cerebellum, and potentially even outside of the cerebellum. Therefore, we tested the generality of the temporally decomposed granule unit basis set.

Using the STP granule unit basis set, we successfully decoded multiple kinematic variables from a motor cortex population response – horizontal and vertical position, velocity, and acceleration of the cursor by fitting weights from granule units to output units (a.k.a. Purkinje cells) (Fig. 6d-f). We used the average firing rates from a publicly-available dataset of single- and multi-unit activity during an isometric wrist force production task^45,46^ (Fig. 6a) as mossy fiber inputs (25 examples in Fig. 6b). We focused on two target directions (+45 and −45 degrees) and modeled 1,000 granule units, where each unit receives inputs from four randomly-selected motor cortex units (Fig. 6c). The same approach decoded EMG from two extensor muscles recorded during the task (Fig. 6g). Crucially, we constrained the decoding so that weights were identical for the +45- and −45-degree directions, though we allowed different weights for different output variables. Thus, the same computation that accounts for the temporal input-output transformation in the floccular complex can transform motor cortical inputs into commands for limb control. Extended Data Fig. 8 shows that the same modeling approach can reconstruct kinematic variables from motor cortex activity during overt center-out reaching movements in primates^47,48^. We do not clai006D that the cerebellum performs this exact set of transformations of motor cortical outputs, but rather that the STP basis set enables multiple transformations even for inputs and outputs that are quite different from those measured in the floccular complex.

**Fig. 6.**
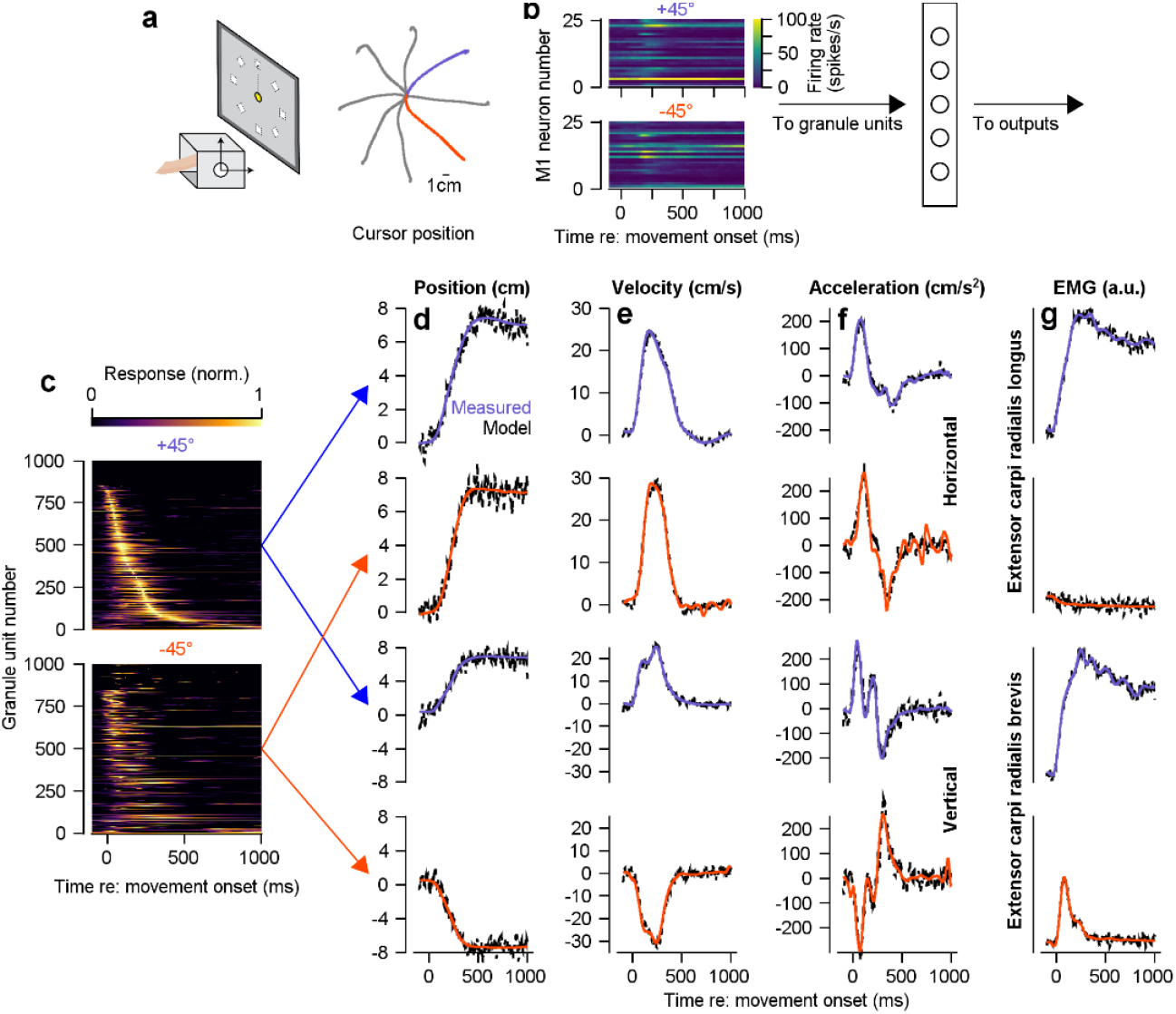
Generalization of the granule unit temporal basis set model to transform the population dynamics in primary motor cortex into the temporal dynamics of cursor kinematics and EMG during isometric force generation in multiple directions. **a**, Schematic diagram of the task (left) and polar representation of force trajectories in 8 directions with the +45 and −45 directions in red and blue (right). **b**, Instantiation of the same STP model as in Fig. 3 but with responses of primary motor cortex neurons as mossy fibers, a layer of granule units, and outputs. The heatmaps show 25 randomly-chosen motor cortex units out of *n* = 3,213. **c**, Heatmaps showing the responses of all 1,000 granule units as a function of time during isometric force generation to targets in the +45 or −45 directions. **d-f**, Cursor position (**d**), velocity (**e**), and acceleration (**f**) as a function of time. Top two rows show horizontal movement, bottom two rows show vertical movement. First and third rows show movement in +45 direction, second and fourth rows show movement in −45 direction. **g**, EMG activity across time measured from two extensor muscles (carpi radialis longus and brevis) during force generation to both targets. Colored and dashed traces show data and predictions of the model. Red and blue arrows between **c** and **d** indicate the granule unit basis sets used to predict the data in each row of **d-f**.

### Links between complex spikes and Purkinje cell directional and temporal tuning

Finally, we ask how the cerebellum might establish, through experience-dependent-plasticity, the appropriate granule cell input weights to create the temporal and directional Purkinje cell response properties we measured. Given that climbing fibers are known to induce plasticity, could the simple-spike responses we measured in floccular Purkinje cells during pursuit have emerged as a consequence of the tuning of climbing fibers? For example, an elegant previous study demonstrated that genetically altering the projections from the inferior olive to the flocculus in mice causes a developmental change in the directional modulation of Purkinje cell simple spikes, dictated by the directional preference of climbing fibers^49^.

Complex-spike firing could dictate simple-spike direction tuning during baseline pursuit because simple-spike and complex-spike responses were anti-correlated, both in absolute preferred direction^32–34,50,51^ (Fig. 7a) and in magnitude (Fig. 7b, R^2^ = 0.16, t(196) = −6.2, p < 10^−9^). In the CS-on vs. CS-off directions, higher vs. lower probabilities of complex spikes were associated with larger decreases vs. increases of simple-spike firing during pursuit.

**Fig. 7.**
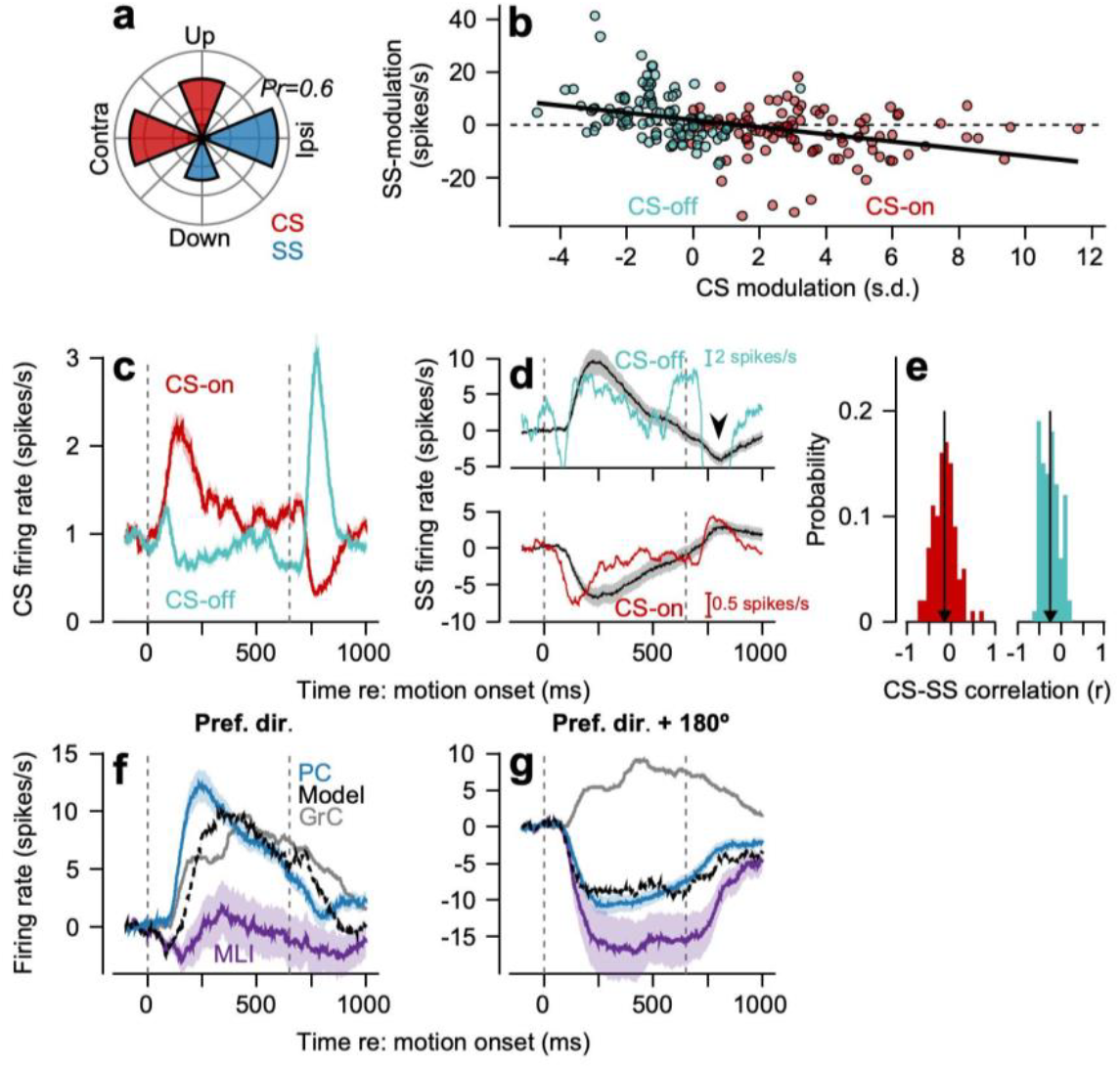
Evidence that complex spike mediated plasticity may create and/or maintain aspects of the temporal and directional dynamics of Purkinje cell firing. **a**, Distribution of preferred pursuit directions for Purkinje cell simple spikes (blue) and their associated complex spikes (red). **b**, Relationship between simple-spike modulation during pursuit in the CS-on (preferred CS) and CS-off (preferred CS+180°) directions as a function of complex-spike modulation across all Purkinje cells. Black line denotes best linear fit. **c**, Red and cyan traces show firing rate of complex spikes in the preferred and anti-preferred complex-spike direction, smoothed using a 50 ms boxcar filter and averaged across Purkinje cells. **d**, Black and colored traces show Purkinje cell simple-spike modulation versus scaled and inverted complex-spike modulation in the CS-off (top) and CS-on (bottom) directions. Arrowhead denotes a region of particularly poor fit between simple-spike and complex-spike modulation. **e**, Histogram of Pearson’s correlation coefficient between the timeseries of complex-spike activity and simple-spike activity in individual Purkinje cells in the CS-on and CS-off directions. Vertical lines denote the population mean. **f**, Simple-spike firing predicted by a model that incorporates the granule unit temporal basis set, complex-spike linked plasticity, and molecular layer interneuron inputs to create the mean Purkinje cell firing rates (blue trace) in the preferred simple-spike direction. **g**, Same **as f** for the anti-preferred simple-spike direction. Note that the simple spike traces differ between panels **d** and **f-g** due to the alignment direction (CS-on versus simple-spike preferred direction). Shaded regions in all panels denote mean ± SEM across neurons.

Alone, the temporal dynamics of complex-spike firing (Fig. 7c) align poorly with those of simple-spike firing during pursuit. In the CS-on direction, the inverse of complex-spike firing starts earlier, rises faster, and decays more quickly compared to simple-spike firing (black vs. red traces, bottom graph of Fig. 7d). In the CS-off direction, the inverse of complex-spike firing shows both an early transient increase and a positive spike after the end of target motion (cyan trace, Fig. 7c), neither of which has a correlate in simple-spike firing (compare black vs. cyan traces, top graph Fig. 7d). Thus, the linear relationship between the timeseries of complex-spike activity to modulation of Purkinje cell simple-spike responses across our population showed negative but small correlations (Fig. 7e, CS-on direction, r = −0.14 ± 0.03; t(99) = −5.4, p < 10^−5^; CS-off direction, r = −0.24 ± 0.02; t(99) = −11.3, p < 10^−18^).

A strong relationship emerged between the timeseries of simple and complex spikes when we considered both the modulation provided by molecular layer interneurons and the properties of the granule unit temporal basis set. We used the successful circuit model of Fig. 4 to adapt the parallel fiber to Purkinje cell synapses on the basis the complex-spike responses of our population of Purkinje cells during pursuit. To obtain the predictions in Fig. 7f,g, we decreased the weights of individual granule unit inputs to Purkinje cells based on the product of each granule unit’s response during pursuit and the mean complex-spike response measured in the same direction; in effect, we decreased weights in relation to the activity of each parallel fiber at the time of a complex spike response. From the excitatory input of granule cells, we subtracted the weighted input of molecular layer interneurons.

The excellent match between observed and predicted mean simple-spike firing in Fig. 7f-g implies that climbing-fiber inputs, when coupled with granule layer activity, could plausibly play a role in creating the temporal dynamics of cerebellar output for the preferred and non-preferred directions. Perhaps there is similar primacy for climbing fiber inputs throughout the cerebellum.

## Discussion

The transformation of neural population dynamics is a ubiquitous problem across brain circuits. Our study identified potential circuit mechanisms in one part of the brain – the cerebellar floccular complex – that could enable two simultaneous transformations along the orthogonal axes of space and time. We suggest that the transformation of the input’s eye position/velocity population dynamics into the output’s eye velocity/acceleration dynamics occurs through (1) decomposition of the incoming signals into a temporal basis set in the structure’s input layer followed by (2) flexible reconstruction, potentially under the guidance of climbing fiber inputs, into any temporal pattern of Purkinje cell firing output. We suggest that the granule cells also reorganize the representation of direction in their mossy fiber inputs, and then excitatory and inhibitory inputs to Purkinje cells can reconstruct a more varied representation of direction in the outputs. The computations in the cerebellar cortex seem to be set up to be incredibly flexible and therefore can reproduce many combinations of directional tuning and temporal dynamics.

Our results are based on one specific cerebellar region, but the ubiquity of the cerebellar circuit suggests that the computational strategies we uncovered have the potential to apply to many behaviors across the cerebellum and potentially throughout the brain. The floccular complex is ideal for our analysis because it provides context that gives strong interpretational power to our results. It is causally required for control of smooth pursuit eye movements^17^, it is connected disynaptically to extraocular motor neurons^19^, and prior recordings had characterized its input-output transformations^21,24^. We have taken the next step by recording the responses of the interneurons that nominally perform the input-output transformations, using new technology to identify neuron types in the cerebellar cortex from extracellular recordings^29,30^, and inferring from both data and computation how the circuit could compute the transformations.

Based on the data, we concluded that the requisite temporal transformation probably occurs in the input layer. In a shorthand, we can think of the mossy fiber inputs to the floccular complex as signaling mostly eye position: identified downstream neurons, including molecular layer interneurons and Purkinje cells^52^, show a temporal transformation to predominantly eye velocity and acceleration. The presence of velocity signals in both molecular layer interneurons and Purkinje cells argues for a shared upstream transformation available to both populations in parallel, rather than independent intrinsic mechanisms reimplemented in each cell type. Golgi cells were essentially unresponsive and likely provide only tonic inhibition to granule cells. Thus, only mossy fibers and unipolar brush cells retain the dynamics of the input signals, meaning that the transformation must be performed through granule cells and/or their synapses on downstream neurons.

We circumvented the challenge that granule cells are inaccessible to conventional recording technologies^29^ by using computational modeling to predict the requisite properties of granule cell dynamics. Myriad analyses revealed that a temporal decomposition of the input signals^53^ by the granule cell population provides an ideal basis set for reconstructing Purkinje cell firing rates. In our system, the basis set must comprise: (1) a broad range of temporal filter responses such that granule cells temporally tile a complete trial; (2) strongly correlated onset and offset time constants resulting in scalar variability^54,55^; (3) granule cell activity that increased with increasing pursuit speeds; and (4) consistent timing of individual granule cell activations across pursuit speed to enable generalization of learning.

We obtained an excellent basis set, good fits to Purkinje cell temporal dynamics, excellent generalization of learning for target speed, and realistic learned timing using biologically motivated short-term plasticity at the mossy fiber to granule cell synapse. Alternative basis sets (see Extended Data Figures) also can support accurate fits, provided the optimization procedure is allowed to select a small subset of units that exhibit the requisite properties. While many parallel fiber to Purkinje cell synapses are likely to be silent, the selection of a diminutive set of “winning” parallel fibers with outsized influence on Purkinje cell dynamics seems non-biological^56^. For this reason, we used an LTP/LTD learning rule to determine parallel fiber-to-Purkinje cell weights, a strategy that distributes weights broadly across granule units.

An engineer might solve the temporal transformation problem differently. For biology, a non-intuitive strategy of decomposing continuous time-varying input signals into a temporally distributed representation and then reconstructing a continuous, but temporally transformed, output has the advantage that it enables a broad range of transformations. It also affords temporally specific learning of sensory-motor input-output transformations^41^. We note that temporal decomposition solutions have been proposed in multiple other papers^40,41,44,53,57^ but based on theory rather than on the neural data we used to come to a tightly constrained solution. We think it is a strength that the properties of our proposed basis set align with features demonstrated as computationally desirable in prior theoretical work.

We also note that prior reports of granule cell responses using calcium imaging are difficult to relate to our hypothesis because (1) the temporal resolution of imaging is limited relative to extracellular neurophysiology and (2) most experimental paradigms did not explicitly establish the causal behavioral role of the recorded regions or quantitatively constrain the relevant behavior. Nevertheless, the relatively dense coding of sensory stimuli in these datasets^58–60^, along with heterogeneous responses across granule cells, would be consistent with formation of a distributed basis set like the one we propose.

Future research will need to test our hypothesis about the nature of the granule cell basis set, explore whether new data suggest different ways to accomplish the same transformations, and evaluate the requisite circuit, cellular, or synaptic mechanisms. Alternatives for the temporal transformation that include downstream cellular mechanisms seem unlikely: (1) parallel fiber transmission to Purkinje cells appears linear up to ~1,000 spikes/second^56,61^; (2) grouped activity in granule cells results in a linear relationship between Purkinje cell response and parallel fiber burst duration at frequencies up to 300 spikes/second^62^, demonstrating that temporal transformations are unlikely to be due to Purkinje cell intrinsic mechanism; (3) Purkinje cells in the vestibulocerebellum show negligible spike frequency adaptation, one plausible mechanism of differentiation^63^.

In the spatial domain, we suggest a homologous transformation of directional tuning in the granule cell layer, one that broadens the directional tunings available to Purkinje cells in their parallel fiber inputs and affords greater flexibility in creating the outputs. Mossy fiber inputs to the floccular complex prefer movements in the cardinal directions – left, right, up, down – while Purkinje cell outputs prefer “ipsiversive” (towards the side of recording) or downward plus slightly contraversive eye motion^22^. Our analysis of the direction tuning in different neuron types implies that parallel fibers with varied multi-direction tunings cross the dendrites of individual Purkinje cells. Multi-directional signals are necessary in the parallel fiber inputs to each Purkinje cell, both to create the various directionality of different Purkinje cells and to enable directional learning in pursuit^42^.

Molecular layer interneurons play a key role in the directional computation, both during baseline pursuit and learning. For baseline pursuit responses, they inhibit Purkinje cells; we suggest that their increases in firing rate for eye movement in their neighboring Purkinje cell’s non-preferred direction drives the Purkinje cell simple-spike responses below their spontaneous firing rates. During learning, molecular layer interneurons provide necessary inhibition that allows a directional depression of simple-spike firing^33,34,64,65^ through complex-spike mediated synaptic depression of directional parallel fiber to Purkinje cell synapses. Thus, the directional transformation in the floccular complex appears to culminate in the molecular layer but still requires combination of different directional inputs in the granule cells.

We confirmed the veracity of the proposed temporal and spatial computations in the floccular complex through simulation of a complete circuit model that predicts the responses of each individual Purkinje cell in our sample. Further, the model has multiple emergent properties that agree with known experimental observations^42,43^ and it performs transformations of temporal dynamics for other datasets from other brain areas.

Fig. 8 schematizes our current understanding of cerebellar computations embedded within the neural circuits that execute pursuit eye movements. The brainstem context is critical because downstream processing constrains what the floccular complex must compute: floccular and vestibular inputs to the brainstem must be temporally integrated to convert velocity commands into the position-dominated signals required by motoneurons^20^. The diverse temporal properties across the basis set enable parallel fiber to Purkinje cell weightings that can reproduce the dynamic response properties of the full sample of Purkinje cells. The diversity of direction tunings allows the system to reproduce the various direction tunings of different Purkinje cells. And the combination of vertical and horizontal direction responses in individual granule cells enables directional learning in pursuit.

**Fig. 8.**
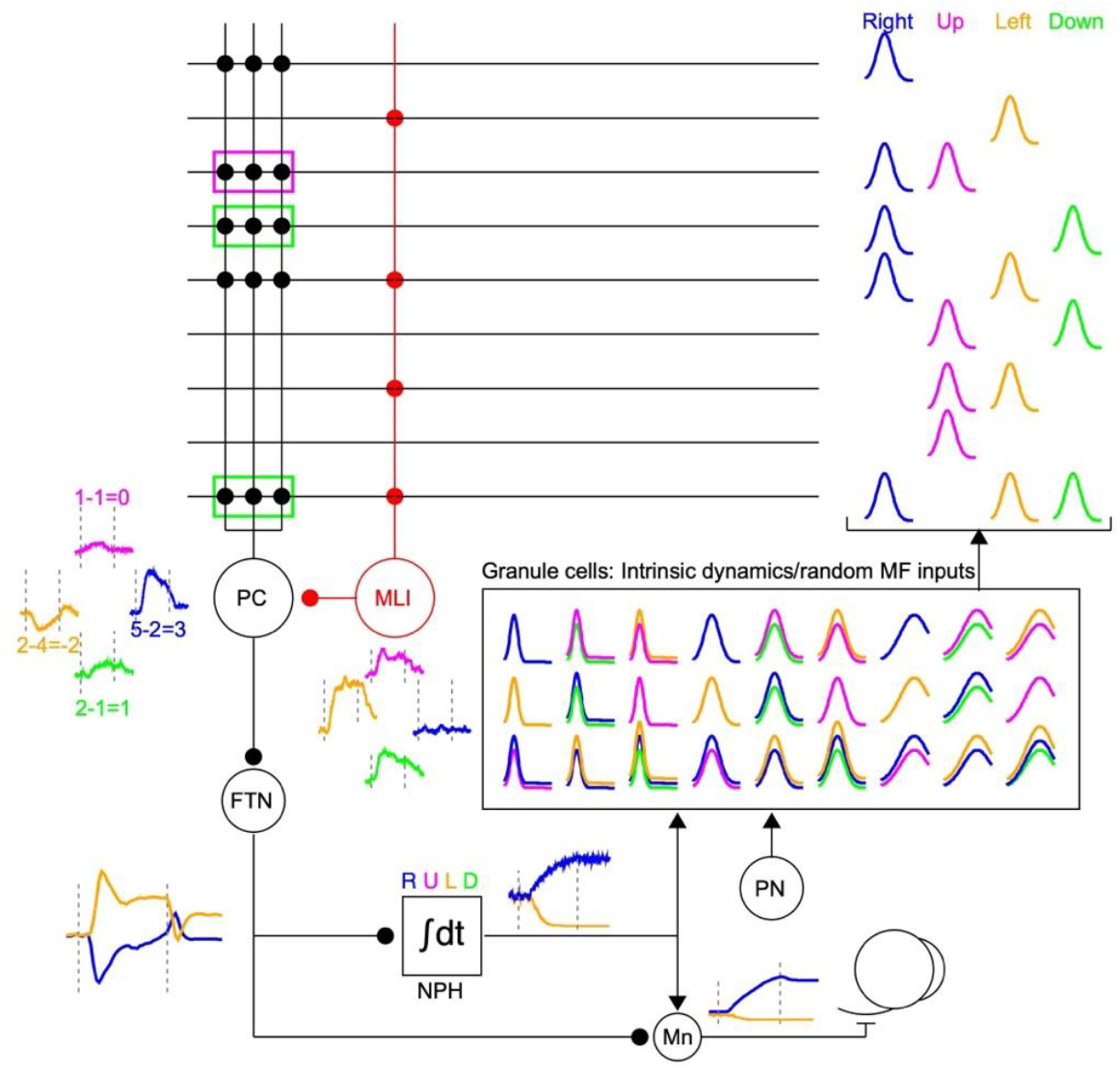
Schematic of the temporal and directional transformations in the floccular complex detailing how they enable temporally-specific directional learning in pursuit eye movements. Throughout the figure, blue, magenta, orange, and green represent neural responses during pursuit to the right (i.e., ipsiversive), up, left (i.e., contraversive), and down, respectively. The brainstem circuits (bottom of figure) illustrate why the floccular complex must transform temporal dynamics to accurately control pursuit eye movements. Eye velocity signals from floccular complex Purkinje cells (PCs) are relayed to floccular target neurons^20,71^ (FTNs) in the vestibular nucleus. Velocity signals are combined with inputs from the vestibular system (not shown) and subsequently integrated (∫ dt) in the nucleus prepositus^20^ (NPH) to create the eye position dominated signal needed by motoneurons^20,72,73^ (Mn). The pontine nuclei^74^ (PN) relay eye signals to the cerebellum via mossy fibers. In the cerebellar cortex, an example basis set with 27 exemplar granule cells spans 3 temporal profiles and 9 combinations of directional tuning (boxed region). In the cerebellum, the granule cells with medium time properties are distributed in a matrix on the upper right that has a row for each parallel fiber and a column for each directional response, showing the distribution of directional preferences for each input. Granule units are combined via weighted synapses onto PCs and molecular layer interneurons (MLIs). The red and black circular symbols show highly-weighted synapses from parallel fibers and the small traces next to the PCs and MLI show the direction tuning expected given these weighted inputs. The numbers on the Purkinje cell traces are the number of weighted parallel fiber-to-Purkinje cell minus the number of parallel fiber inputs supplied to the connected MLIs. The magenta and green rectangles on parallel fiber to Purkinje cell synapses show those that have an appropriate directional signal to allow pursuit learning for a leftward instruction presented during upward and downward pursuit, respectively. The different time courses of the 3 groups of granule cell responses allow the system to reproduce the varied time courses of firing rate in different Purkinje cells and, after Buonomano and Mauk^41^, enable temporally-specific learning.

Our analysis provides a roadmap for asking whether the cerebellar cortex performs a “universal transform” that aligns with the universal architecture of the cerebellar circuit^13,14^. For example, existing data in the eye-blink regions in mice^66^ and rabbits^67^ suggests a temporal basis set analogous to that proposed here for smooth pursuit, reinforcing the idea that similar circuit computations may generalize across cerebellar behaviors. And, as we showed for the responses of neurons in motor cortex, the temporal basis set provided by our model granule units could support many different temporal transformations on the time scale of movements. Indeed, the principle of combining varied inputs onto granule cells to create a basis set with a wide range of representations of cerebellar inputs could extend beyond time, for example to direction in our model. It would maximize the flexibility in learning of appropriate cerebellar output to guide behavior. Multiple plasticity mechanisms^68,69^, including meta-plasticity^70^, could further generalize the framework of a high-dimensional basis set to the specific demands of different behaviors and cerebellar areas.

In summary, we describe, for the first time in a behaving vertebrate, how a complete circuit with input, output, and interneurons could perform specific neural computations. Our analysis points to biologically plausible neural circuit mechanisms that can transform neural population dynamics across space and time and suggests that these circuit mechanisms could be a fundamental feature of neural circuits throughout the cerebellum and the rest of the brain.

## Methods

All experimental procedures were approved in advance by the Duke *Institutional Care and Use Committee* under protocols A085-18-04, A062-21-03, and A016-24-01. All animal care and experimental procedures followed the guidelines outlined in the *NIH Guide for the Care and Use of Laboratory Animals* (1997). Three rhesus macaques (*Macaca mulatta*, all male, 10-15 kg) were used in this study. Portions of the dataset analyzed for this manuscript have been included in three previous reports^29,30,32^.

### General procedures

Prior to behavioral training or neurophysiological recordings, monkeys were instrumented with a head-restraint device, an eye coil, and at least one recording cylinder via separate surgical procedures, all performed with aseptic technique. For each surgical procedure, animals were deeply anaesthetized with isoflurane. Animals received peri- and post-operative analgesia until they had completely recovered. In the first of multiple surgical procedures, head restraint hardware was attached to the animal’s skull to allow measurement of the animal’s eye position without concomitant head movements. In a later surgery, we sutured a small coil of wire to the sclera of one eye^75^, allowing measurement of the animal’s eye position using the search coil technique^76^. Following these two surgical procedures, animals were trained to perform smooth pursuit eye movements during discrete trials of target motion in exchange for fluid reward. The experimental paradigm is described in detail below. Once the animal had demonstrated excellent smooth pursuit tracking abilities, as evidenced by minimal intervening saccadic eye movements, we affixed a stainless-steel recording cylinder above a craniectomy, allowing access to the floccular complex of the cerebellum with microelectrodes. The position of this cylinder was 11 mm lateral to the midline, angled at 26° relative to the frontal plane, and pointed towards the interaural axis.

### Behavioral task

Each day animals were seated in a dark room, 30 cm in front of a CRT monitor (80 Hz refresh rate, 2304 × 1440 pixels, 480 × 310 mm). Their heads were attached to the chair via the previously implanted restraint hardware. Horizontal and vertical eye position signals from the search coil system were separately digitized at 1,000 Hz. We computed the velocity of the animal’s eye movement offline using a 2^nd^ order causal Butterworth low-pass filter with a cutoff frequency of 10 Hz. As we were principally interested in the relationship between floccular neural responses and smooth pursuit eye movements, we removed any saccadic eye movements from ongoing pursuit using an automated procedure^32^. The occurrence of a saccade was identified offline using eye velocity (50 deg/s) and eye acceleration thresholds (1,250 deg/s^2^). Onset and offset of each saccade were determined as the time when the animal’s eye velocity or acceleration fell below both thresholds for more than 10 ms. In all analyses, we treated eye kinematics during saccades as missing data.

Stimulus presentation was controlled via our lab’s custom “Maestro” software. The visual stimulus in all trials was a small (0.5° diameter) black spot shown on a light gray background. In a small subset of trials, animals were required to fixate a stationary target placed at one of nine locations, evenly spaced within a 10° x 10° square. The monkey was required to fixate the dot at each position (within ±1°) continuously for one second in exchange for reward. In the majority of trials, animals tracked smoothly moving targets. Each smooth pursuit trial began with the monkey fixating the dot in the center of the screen for a randomly chosen intertrial interval (400-800 ms, uniformly distributed). After the fixation interval, we used Rashbass’ step-ramp paradigm^77^. On each trial, the target moved in the “pursuit direction” with a speed of 10, 20, or 30 deg/s. At the onset of target motion, the target was stepped backwards by 1.5°, 3°, or 4.5° for each of the three target velocities. The backwards step minimizes catch-up saccades caused by the visual latency of the pursuit system^78^. The target moved at constant velocity for 650 ms before stopping at an eccentric position for an additional 350 ms. The majority of pursuit trials were performed in the four cardinal directions. Animals were rewarded for keeping their eyes within an invisible 3° bounding box centered on the pursuit target for the duration of the trial, including initial fixation, pursuit, and eccentric fixation at the end of the trial. Animals were trained extensively on the pursuit task prior to neurophysiological recordings.

### Neurophysiology recordings

All recordings were made in the ventral paraflocculus and flocculus, a region we call the floccular complex. These regions have been shown to be crucial for the execution of smooth eye movements^17^. In addition, electrical stimulation of this region of the cerebellum drives smooth movement of the eye towards the side of stimulation^20,71^. We acutely inserted either single tungsten microelectrodes (FHC, 1-2 MΩ) or, more commonly, custom designed Plexon S-Probes into the brain each day through the previously implanted recording cylinder. Plexon S-Probes featured 16 tungsten recording contacts, each 7.5 µm in diameter. The 16 contacts were arranged in two columns (50 x 50 µm separation between adjacent contacts). Almost all of our recordings come from the S-Probes.

We identified the floccular complex by its strong activity during smooth pursuit eye movements, presence of infrequent Purkinje cell complex spikes, and the depth relative to the tentorium. Upon arriving in the floccular complex, we waited a minimum of 30 minutes, up to several hours, before beginning neurophysiological recordings. This initial period maximizes the stability of the recording by minimizing drift of neural units across the recording contacts.

Neurophysiological data were recorded using the Plexon Omniplex system. We used analog Butterworth low-pass hardware filters (4^th^ order, 6 kHz cutoff) prior to digitization to minimize any interference from the eye coil system and prevent aliasing. Wideband neural activity on each contact was recorded at 40 kHz, synchronized to behavioral data using high speed TTL pulses, and stored for later offline analysis.

To identify well-isolated single neurons from our wideband voltage recordings, we leveraged the Full Binary Pursuit^79^ (FBP) spike sorting package. The FBP sorter is specifically designed to optimally resolve temporally and spatially overlapping action potentials, a frequent occurrence in the cerebellar cortex due to the relatively high baseline firing rates of many cerebellar neurons. Following spike sorting, we manually curated all neural units to ensure high quality single-units. We specifically excluded neural units that showed evidence of contamination by either background noise or other units. We assayed contamination primarily via assessment of refractory period violations, measured as the percentage of spikes that violated a presumed 1 ms absolute refractory period. Across all monkeys, we recorded *n=1,152* single units. Mean refractory period violations were 0.6 ± 2.5% (mean ± SD) of all recorded spikes. We measured the maximum peak-to-peak voltage deviation of the mean action potential waveform/template on the channel with the largest spike and compared this amplitude to the level of background noise measured on the same channel (the primary contact). The mean signal-to-noise ratio, defined as the amplitude of the spike divided by an estimate of noise computed as 1.96-times the standard deviation of background noise, was 5.6 ± 2.9 (mean ± SD). We did not consider the functional properties of neural units when manually curating output of the sorter, opting to analyze all neurons recorded from the floccular complex with acceptable isolation. We converted neuron spike trains into firing rates by causally convolving each spike train with a dual exponential kernel. The kernel (τ_rise_ = 0.1 ms and τ_decay_ = 50 ms) mimics the properties of post-synaptic currents^80^ while preserving the ability to measure latencies with high precision. Autocorrelograms were computed using previously described techniques^32^, and normalized by the bin width (1 ms), placing autocorrelograms in units of spikes/s.

### Expert identification of cerebellar neuron types

A detailed discussion of the methodology for identification of cerebellar neuron types in our recordings has been published previously^30^. Briefly, we began by identifying a subset of units as ground-truth Purkinje cells based on their unique extracellular properties. Purkinje cells receive excitatory inputs from parallel fibers as well as strong inputs via climbing fibers from the inferior olive that drive post-synaptic Purkinje cell complex spikes. During complex spikes and for 10 or more milliseconds thereafter, Purkinje cells do not fire simple spikes, resulting in a stereotypical complex-spike-induced pause in simple spikes. We included only Purkinje cells that had a complex-spike-induced pause.

We identified molecular layer interneurons by establishing monosynaptic functional inhibitory relationships between these neurons and simultaneously recorded ground-truth Purkinje cells. Therefore, our sample of molecular layer interneurons constitutes principally MLI-1’s, classified by others based on their genetic^25^ and connection profiles^81^. Golgi cells were identified by their low firing rate responses, characteristic extracellular waveforms^82^, and presence within the granule cell layer. Mossy fiber inputs to the cerebellar cortex were identified based on the presence of a negative-after-wave^83^ (NAW). The NAW corresponds to the postsynaptic response of granule cells within the cerebellar glomerulus^84^. We note that while a NAW is sufficient to identify mossy fibers^29^, it is possible to record from mossy fiber axons that do not show marked NAWs. To ensure that our sample contained only known mossy fibers, we excluded from analysis putative mossy fibers that lacked a NAW. Unipolar brush cells were identified based on established functional responses recorded by *in vitro* studies showing that unipolar brush cells elongate the timescales of discrete mossy fiber input over 10s to hundreds of milliseconds^85,86^. We considered units to be unipolar brush cells if triggering their firing off simultaneously recorded mossy fiber bursts yielded temporally-elongated responses. We validated our expert-labeling approach using a deep-learning classifier trained on ground-truth optogenetically-identified recordings in mice^29^.

### Principal component analysis

To identify the primary modes of temporal information contained in cerebellar populations during smooth pursuit eye movements, we performed principal component analysis across all expert-identified neurons. As the responsiveness of cerebellar neurons to pursuit varied widely, we wanted to ensure that highly responsive neurons did not bias our estimate of the principal component directions. Therefore, we performed soft-normalization of neuron firing rates^31^, measured in their preferred direction, prior to principal component analysis. Soft-normalization squashes modulations less than 5 spikes/s to timeseries near zero and scales modulations of greater than 5 spikes/s to approximately unit magnitude. After soft-normalization, we smoothed the firing rate traces using a boxcar filter with a 50 ms width. To identify the principal components across speed and direction for mossy fibers and Purkinje cells, we used the same preprocessing procedures separately in each respective population. Then, we concatenated the within-neuron responses to different speeds and directions, resulting in an N x (S*D*N_t_) matrix, where N represents the number of units in each population, S represents the number of speeds (three: 10, 20, 30 deg/s), D the number of directions (four: ipsiversive, up, contraversive, down), and N_t_ the number of timepoints in each sample (1,100). As with the complete population, principal components analysis identified the modes of temporal profiles across the neuron dimension in each population.

### Regression of neuron firing rates to eye kinematics

To evaluate the contributions of different eye kinematic signals to the firing rates of cerebellar neuron types, we performed linear regression analysis by fitting kinematic models to each neuron’s mean time-varying firing rate during smooth pursuit. Depending on whether the firing in the anti-preferred direction hit a floor at zero firing rate, we fit the data either in the neuron’s preferred direction or across both its preferred and anti-preferred directions simultaneously. For each neuron, we fit models of the following form:

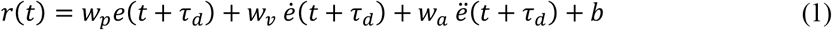

In Equation 1, *r(t)* represents the mean firing rate of the neuron across pursuit trials and is expressed as a weighted sum of the mean eye position (*e*), velocity (ė), and acceleration (ë) at a future timepoint, (*t* + *τ*_*d*_). The parameter *τ*_*d*_ refers to the temporal lead of the firing rate relative to the kinematics. The scalar parameter, *b*, is a neuron-specific bias. We determined the unknown parameters, {*w*_*p*_, *w*_*v*_, *w*_*a*_, *b*} using least squares for each delay from 0 to 100 ms. The optimal delay was identified by minimizing the mean squared error across the range of delays. To assess the relative contribution of each kinematic variable to the overall firing rate, we computed standardized (*β*) coefficients by normalizing each weight by the ratio of the standard deviation of the corresponding regressor to the standard deviation of the observed firing rate.

### Deep-learning neural network model of granule cell transformations

We constructed a neural network model to investigate features of the transformation between mossy fiber and unipolar brush cell inputs and Purkinje cell outputs. We supplied as inputs to the model single-trial firing rate responses of individual mossy fibers (*n = 86*) and unipolar brush cells (*n = 31*) during 10, 20, or 30 deg/s smooth pursuit trials measured in one of the four cardinal directions. The model was trained to predict the single-trial firing responses of our recorded Purkinje cell population in their preferred directions. Input and output trials were trial-matched for speed (e.g., mossy fiber and unipolar brush cell inputs corresponding to single-trial responses during 10 deg/s target motion were paired with Purkinje cell outputs also measured during 10 deg/s pursuit). Choice of the direction of pursuit for mossy fiber and unipolar brush cells is arbitrary, as we noted a relatively uniform distribution of preferred directions of these input neurons across these directions. However, we chose to fit transformations of Purkinje cells only in their preferred direction because it minimizes the influence of molecular layer interneurons, whose activity does not, on average, modulate in simultaneously recorded Purkinje cells’ preferred direction. Thus, fitting specifically Purkinje cell preferred directions enables us to reduce the complexity of the fitted model.

The model consisted of 100 “granule units.” Each granule unit received input from four randomly selected mossy fibers or unipolar brush cells, summed with non-negative, uniformly randomly selected weights between 0 and 1 as described in Equation 2.

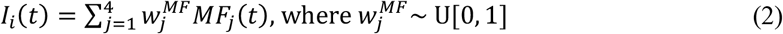

In Equation 2, *MF*_*j*_(*t*) represents the firing rate of either a single mossy fiber or unipolar brush cell (intrinsic mossy fiber).

To capture the dynamics needed to transform inputs into Purkinje cell outputs, each granule unit filtered its input, *I*_*i*_(*t*), through 25 first-order linear filters with shared time constants, **τ**.

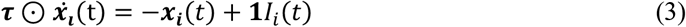

In Equation 3, ***x***_***i***_(*t*) is a 25-element vector of filter states. Each granule unit then combines these filter outputs with unit-specific weights and passes the result through a saturating softplus activation function, σ, to enforce non-negative firing rates according to Equation 4.

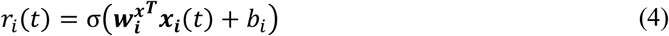

While time constants were shared across all granule units, each unit had its own input connections, filter weights 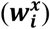, and bias (*b*_*i*_). In this and all subsequent models, we subtracted the bias from the inputs and then fitted the bias as part of the optimization procedure to allow the model to reproduce neuronal firing rate modulation without being constrained by baseline firing rates.

Finally, a set of fully connected weights mapped the *n* = 100 granule units timeseries (*R(t)*) onto each output Purkinje cell: 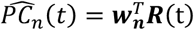. Again, we constrained the weight matrix between granule units and Purkinje cells to be strictly positive to represent the excitatory nature of the mossy fiber to granule cell and parallel fiber to Purkinje cell synapses.

The complete network was simultaneously trained using the Adam^87^ optimizer with cosine annealing with warm-restarts^88^. We used drop-out layers during training with 50% probability of dropout after the granule unit layer to avoid overfitting and ensure that the dynamics necessary to represent Purkinje cell firing rates were distributed throughout the granule layer units. We used early termination to stop training when the cross-validated error from a withheld 10% of the single trial data failed to decrease for more than 10 training epochs.

We evaluated performance of the network by supplying as input the trial-averaged activity (rather than single-trial activity) of mossy fibers and unipolar brush cells and measuring the network predicted Purkinje cell responses. We compared the predicted responses to the mean measured Purkinje cell responses across target speeds. We also examined the nature of the computation performed by the complete network. To do so, we supplied a step-response of mossy fiber and unipolar brush cell activity as input. At *t=0* ms, we stepped the response of each mossy fiber to the baseline-subtracted response measured at the termination of target motion across pursuit in the 20 deg/s condition. We then interrogated the responses of the granule units across time from before to the end of the step change in input.

### Approach to evaluate different models of granule cell dynamics

Given the inability to record cerebellar granule cells via extracellular recordings^29^, we sought an unbiased approach to evaluate different models of granule cell dynamics. We created 5 model granule unit basis sets, one that emulates short term synaptic plasticity and four others suggested in the literature or by reviewers (see below). For each model basis set we optimized the mapping from mossy fiber and unipolar brush cell inputs, as measured in one of the cardinal directions, to Purkinje cell outputs as measured in their preferred direction of pursuit. The logic is the same as described above: focusing on Purkinje cell responses in the preferred direction allows us to reduce the model by eliminating temporal effects of molecular layer interneuron firing.

As we were chiefly interested in the characteristics of the basis sets formed by each model, the parameters of individual granule units were chosen randomly. Each free parameter of the model (described below) was drawn from a bounded uniform distribution whose hyperparameters (lower and upper bounds) were optimized to reproduce Purkinje cell firing rates. We used black-box optimization (adaptive differential evolution) to identify the lower and upper bounds of the uniform distribution describing each free parameter. We minimized the mean squared error between the reconstructed and measured Purkinje cell activity across our complete sample of *n = 101* Purkinje cells. For each iteration of the optimization algorithm, we constructed a random selection of 100 granule units, with each unit’s parameters drawn from their respective distributions. We used an LTP/LTD rule to determine the weight matrix between granule units and Purkinje cells (described below) and averaged the mean squared error across five replicates per optimizer iteration.

For the majority of analyses, we used an iterative procedure to identify the weights between granule units and Purkinje cells. The learning rule is based on the observation that the absence of climbing fiber activity in the cerebellum drives functional potentiation whereas complex spike firing drives depression at the parallel fiber-to-Purkinje cell synapse^33,64,89^. In the *n*-th iteration of the learning algorithm, we computed the Purkinje cell reconstruction 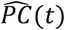 as the weighted sum of granule cell responses:

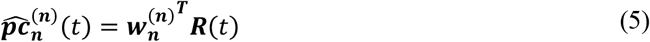

In Equation 5, ***w*** is a vector of granule cell weights and ***R*** is the response of each granule cell. The reconstruction error is the difference between the measured and reconstructed Purkinje cell responses: 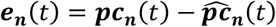. Reconstruction errors greater than zero drive depression whereas errors less than zero drive potentiation, according to Equation 6:

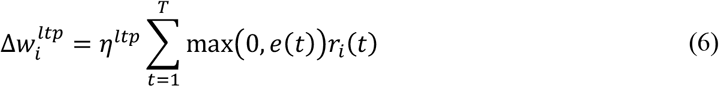

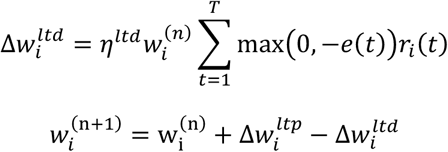

The potentiation term in Equation 6 increases weights for granule cells whose activity is correlated with underestimates in the reconstruction. In contrast, granule cells whose activity is positively correlated with overshoots are downweighted by the depression term. Note that the depression term is multiplicative with the current synaptic weight, 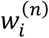, which ensures that weights remain strictly positive. The learning constants η^ltp^ and η^ltd^ are somewhat arbitrary, provided they are sufficiently small to ensure convergence (which depends on the number of granule units; in practice, we scaled these constants by the number of granule units).

The learning rule described in Equation 6 is important for testing the generalization properties of different models of granule cell dynamics. The iterative update defined in Equation 6 spreads weight updates across granule cells whose dynamics correlate with the current reconstruction error. In most models of granule cell dynamics, strong correlations exist between granule units with similar parameters. Alternative methods for determining the weight matrix, such as least-squares approaches, will likely achieve lower mean squared reconstruction error than Equation 6, but yield weight matrices that are difficult to interpret in the context of the cerebellum due to multicollinearity. Thus, in several analyses noted in the main text, we use least-squares approaches to determine whether sufficient information exists in the granule cell basis set to reconstruct specific signals (e.g., Fig. 6 and Extended Data Fig. 8). However, for analyses requiring interpretation of the weight matrices, we use the learning rule described in Equation 6.

### Creation of models of granule unit basis sets

#### A short-term synaptic plasticity model of granule cell activity

As short-term synaptic plasticity at the mossy fiber-to-granule cell synapse has been hypothesized to enrich granule cell dynamics^44^, we developed a model to test whether short-term plasticity could establish a granule cell basis set capable of recapitulating our Purkinje cell firing responses and generalize to novel learned contexts. Prior experimental^36,37^ and computational^44^ work provides the basis for this model. As in the neural network model, we modeled the input to each granule unit by selecting four random mossy fibers and unipolar brush cells as presynaptic inputs. Each synapse was subject to short-term plasticity using an extension of the classic Tsodyks-Markram framework^38^. Briefly, each synapse features two pools of synaptic transmitter. The first pool is fast and readily releasable, while the second pool is slow and serves to replenish the first. Because the Tsodyks-Markram model operates at the level of individual action potentials, we converted presynaptic mossy fiber and unipolar brush cell firing rates into the probability of observing a spike at each time step:

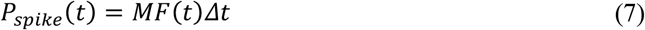

As in Equation 2, *MF(t)*, represents the firing rate of mossy fibers or unipolar brush cells relative to baseline. Multiplication by the time step Δ*t* (1 ms), convert firing rates to spiking probability per millisecond bin.

We model synaptic facilitation using the classic Tsodyks-Markram formulation, governed by the differential equation in Equation 8:

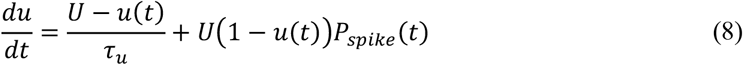

In Equation 8, *U* is the baseline utilization and *τ*_*u*_ is the facilitation timeconstant. The current synaptic utilization *u(t)* is clamped between zero and one. We drew *τ*_*u*_ for each synapse from a uniform distribution between 2 and 10 ms, consistent with observed rapid facilitation at this synapse (e.g., ~12 ms observed by ref. ^37^).

Release of synaptic transmitter is determined by the current availability in the fast pool *R*_*fast*_(*t*), the probability of a presynaptic action potential, and the current fractional utilization, via Equation 9.

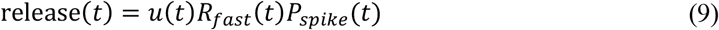

Neurotransmitter release from the fast pool reduces the amount available:

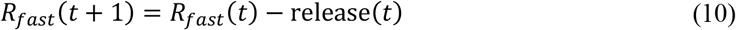

The fast pool is gradually replenished by transfer from the slow pool, with recovery time constant *τ*_*fast*_, described in Equation 11. This transfer adds transmitter to the fast pool while reducing the slow pool:

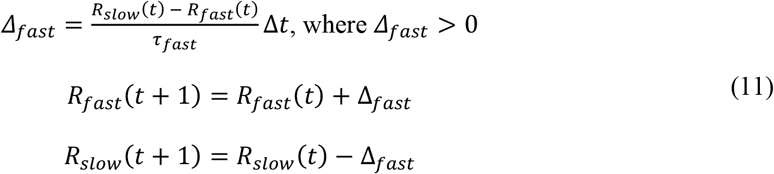

The slow pool then recovers with time constant *τ*_*slow*_, according to Equation 12.

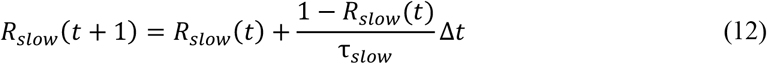

The total synaptic current equals the released neurotransmitter multiplied by an arbitrary scalar synaptic weight drawn from a uniform distribution between zero and one: *I*_*j*_(*t*) = *w*_*j*_ release(*t*). The final granule cell firing rate is given by Equation 13:

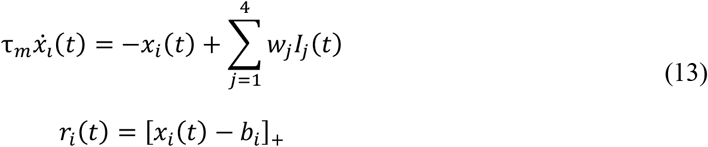

In Equation 13, *τ*_*m*_ is the granule cell membrane time constant, drawn from a uniform distribution between 1 and 5 milliseconds. The firing rate of the granule unit, ***r***_*i*_ is constrained to be non-negative by half-wave rectification, denoted by [·]_+_. Thus, this short-term synaptic plasticity model had four free parameter distributions governing the dynamics: the bias *b*, the baseline synaptic efficacy *U*, and the 2 time constants, *τ*_*slow*_, and *τ*_*fast*_.

#### Alternative models of granule cell dynamics

We tested multiple alternative granule cell models to determine the set of properties necessary to reproduce Purkinje cell firing rates and generalize across learning contexts. Below, we describe four alternative models: (1) a passive model that transmits combined mossy fiber information, (2) a model with a compressive nonlinearity, (3) a model with activity-dependent thresholds, and (4) a model that uses recurrent activity to produce a basis set. As with the short-term plasticity model (Equations 7–13), we optimized the hyperparameters of the distributions governing the free parameters to minimize the squared error between reconstructed and measured Purkinje cell activity in each cell’s preferred direction using the LTP/LTD plasticity rules in Equations 5 and 6. Note that all models allow optimization of the bias of granule units, enabling them to be tuned to move all sustained inputs below threshold and create transient responses that tread on the noise in their responses.

Across all models, each granule unit received input from four randomly selected mossy fibers and unipolar brush cells with random weights (uniformly distributed between 0 and 1), as described in Equation 2. In the passive model, granule units performed no processing beyond the application of a bias and half-wave rectification. The response of the *i*-th granule unit was therefore given by Equation 14:

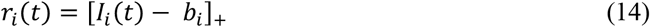

The notation in Equation 14 is identical to that in Equation 13: *b*_*i*_ is a unit-specific bias drawn from an optimized uniform distribution, and the brackets denote half-wave rectification at zero. The passive model had a single free parameter distribution, corresponding to the bias.

In the compressive nonlinearity model, we used a sigmoid activation function as an exemplar compressive nonlinearity. The activity of each granule unit in the saturating nonlinear model is defined in Equation 15:

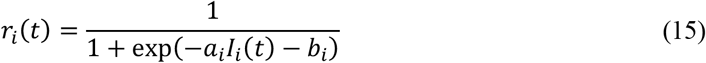

In the model described in Equation 15, two parameter distributions were optimized: the unit-specific gain *a*_*i*_ and the unit-specific bias *b*_*i*_.

In the thresholded granule unit model, incoming mossy fiber activity was subject to a unit-specific bias, as in Equation 14. However, rather than drawing the bias from an optimized uniform distribution, the threshold was derived from the activity profile of the input using a previously proposed formulation^90^:

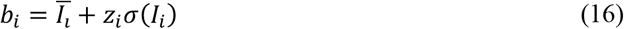

In Equation 16, Ī_*i*_ represents the mean input across time and pursuit conditions and *σ*(*u*_*n*_) represents the standard deviation of the input across time and pursuit conditions. In essence, Equation 16 sets the threshold based on the z-scored input to each granule unit. The condition where *z*_*i*_ = 1 corresponds to a granule unit whose input does not exceed the threshold for activation 50% of the time. This model had one parameter distribution that we optimized, corresponding to the z-score threshold, *z*.

In the random projection model, granule cell activity is shaped by recurrent inhibition from randomly connected units, a model that has been proposed in multiple forms^41,57,91,92^. The activity of granule units in the random projection model is given by Equation 17:

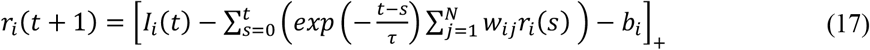

In Equation 17, the activity of the *i*-th granule unit is the difference between the incoming mossy fiber activity and inhibitory inputs from temporally weighted recurrent network activity – typically attributed to input from the activity of Golgi cells. As in Equations 13 and 14, the brackets denote half-wave rectification. The optimized parameter distributions included the time constant τ, governing the exponentially weighted history of recurrent activity, and κ, governing the recurrent connection weights. The recurrent connection matrix was drawn from a binomial distribution such that *Pr*(*w*_*ij*_) = 0 = *Pr*(*w*_*nj*_ = 2κ/*N*) = 0.5. Here, *N* is the total number of granule units in the simulation.

### Cerebellar circuit models for predicting downstream neural responses

Using the short-term synaptic plasticity granule unit basis set, we asked whether the resulting granule unit population could replicate the measured firing rate responses of neural units downstream of granule cells: molecular layer interneurons and Purkinje cells. We constructed a population of 1,000 simulated granule units using the procedures outlined above and computed their predicted firing rate responses to pursuit at 20 deg/s in the cardinal directions.

We first modeled the responses of molecular layer interneurons. Across all cardinal directions simultaneously, we applied the learning rule in Equation 6 to find a set of weights that predicted molecular layer interneuron activity from granule unit activity. Because our population of molecular layer interneurons was relatively small, we augmented it by randomly averaging sets of 5 recorded molecular layer interneurons with replacement. Fits were evaluated using Pearson’s correlation coefficients.

We then evaluated the ability to fit our measured Purkinje cell responses across directions. We applied the learning rule in Equation 6 to simultaneously identify granule cell and molecular layer interneuron weights that predicted individual Purkinje cell responses. Molecular layer interneuron responses were multiplied by −1, so that non-negative weights corresponded to inhibitory inputs onto Purkinje cells, and directionally permuted so that interneuron preferred directions were opposite to Purkinje cell preferred directions, as observed in our recordings. Our goal was therefore to find a set of non-negative weights describing Purkinje cell firing across the measured population, given the joint contributions of upstream granule cell excitation and molecular layer interneuron inhibition.

### Evaluation of emergent properties of the cerebellar circuit models

We tested whether our simplified model of cerebellar function could account for previous pursuit learning results. Previous results suggest that short-term pursuit learning is driven by complex-spike mediated plasticity of the parallel fiber to Purkinje cell synapse. A complex spike on a single learning trial is linked to a well-timed depression of Purkinje cell simple-spike firing on the next trial^33,34,64,65,89^. To simulate single-trial depression in our cerebellar circuit model, we modified the weights between granule cells and Purkinje cells identified by each granule unit model. Each granule cell’s learned weight, *w*_*i*_, is:

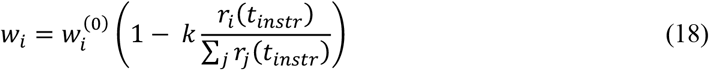

In Equation 18, ***r***_*i*_ is the activity of each of 1,000 granule units measured at the time of the instruction: synapses from inactive granule units would not be depressed. The scalar *k* scales the magnitude of Purkinje cell learning. To enable comparisons across models with widely varying granule unit activity scales, *k* was normalized by the total population activity at the time of the instruction. Learning was quantified as the simulated change in Purkinje cell simple-spike output between the original weight vector 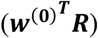 and the modified weight vector (***w***^***T***^***R***) where ***R*** is the matrix of granule unit activity over time.

In one set of simulations, we tested generalization of learning by changing the statistics of the mossy fiber inputs on the test trial after each learning trial, thereby assessing learning for a range of pursuit speeds in the test trial. In a second set of simulations, we changed the timing of the instructive stimulus relative to the onset of pursuit and asked how that altered the Purkinje cell firing on the subsequent test trial. We emphasized generalization of learning in the main paper by using the STP granule unit basis set, but we also performed full circuit optimization with the four alternative granule unit basis sets to assess how well they could support the observed generalization (Extended Data Fig. 3-4). Finally, we repeated the entire extent of our assessment of different granule unit basis sets using non-negative least squares rather than LTP/LTD as the learning rule (Extended Data Fig. 5-7).

### Generalizability of the temporal transformation to limb movements

Using the short-term plasticity model (Equations 6-11), we asked whether this basis set could account for transformations in other regions of the cerebellar cortex, beyond the floccular complex, during different motor control tasks. We analyzed a publicly available dataset^45,46^ of single- and multi-unit activity recorded from the motor cortex of rhesus monkeys performing an isometric wrist force production task toward one of eight “force targets.” Intramuscular EMG activity from multiple muscles was recorded simultaneously with neural activity from a 96-contact Utah array chronically implanted in primary motor cortex. In total, we analyzed activity from 3,213 contacts across multiple sessions in two monkeys.

As above, we constructed a population of 1,000 model granule units, each receiving as input the mean activity of four randomly selected motor cortex contacts (with uniformly distributed random weights between 0 and 1). To test whether the temporal decomposition of motor cortex activity by the granule cell model contained sufficient information to reconstruct kinematic and EMG activity, we decoded multiple potential cerebellar outputs as non-negative weighted sums of granule unit activity. Each putative output, envisioned as a Purkinje cell, was obtained using a single weight matrix fit by non-negative least squares simultaneously across the +45 and –45 degree force targets. This approach not only decoded granule unit activity but also prevented interference across the two directions. To account for biases in the output variables (e.g., EMG signals), we augmented the 1,000 granule units with positive and negative bias terms, also fit via non-negative least squares. We performed an identical analysis on a separate dataset of motor cortex activity (n = 537 units) recorded during overt reaching movements in monkeys^47,48^, although EMG was not available for this task.

### Purkinje cell responses reflect the statistics of granule unit and complex spike activity

Complex-spike-mediated synaptic plasticity is thought to be a critical driver of Purkinje cell activity in the cerebellum. We used computational modeling to test whether the joint activity of granule cells and climbing fiber responses might have created the temporal trajectory of Purkinje cell firing. Specifically, does the measured complex spike activity, combined with granule cell activity predicted by the short-term plasticity model, predict the temporal responses of Purkinje cells? We tested this using a modification of the iterative learning rule in Equation 6. Equation 19 parallels Equation 6, with the difference that rather than splitting the temporal error vector into positive and negative components to drive learning, we used the measured complex spike activity for each Purkinje cell, relative to baseline, as the plasticity signal:

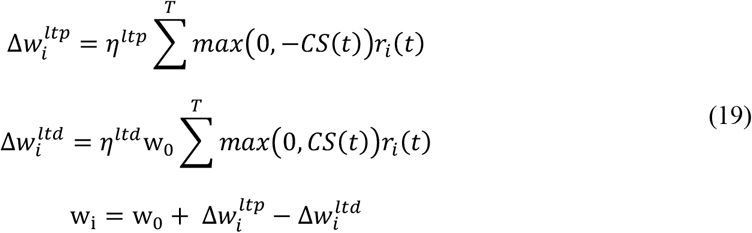

In Equation 19, *CS(t)* represents the mean measured complex spike activity relative to baseline. We then identified optimal scalar weights for granule cell and molecular layer interneuron inputs across all Purkinje cells to reconstruct Purkinje cell responses:

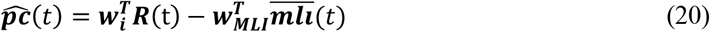

In Equation 20, ***R***(t) represents the input of granule units across time, ***w***_***i***_ from Equation 19 and 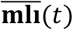 represents the mean molecular layer interneuron activity. Thus, the complete model has 4 free parameters across all Purkinje cells: w_0_, the baseline weight, two plasticity rates (*η*^*ltp*^ and *η*^*ltd*^), and a scalar weight representing the mean input of molecular layer interneurons across Purkinje cells, **w**_**mli**_, which we fit using gradient descent.

### Analysis of spike timing synchrony among mossy fibers

We evaluated the millisecond-scale synchrony between pairs of simultaneously-recorded mossy fibers using techniques we developed to assay Purkinje cell simple-spike synchrony^32^. Briefly, we assayed the probability of the two mossy fibers firing in the same millisecond during a smooth pursuit trial, *Pr*[*MF*_1_(*t*) ∩ *MF*_2_(*t*)]. This raw measure of synchrony is the combination of synchrony that is expected due to temporal covariations in the firing rates of both mossy fibers as well as potential synchrony that is due to network effects independent of rate-based changes. To separate these two potential sources of millisecond-scale synchrony, we subtract from each raw measure of synchrony the expected synchrony under the null hypothesis that the two neurons are independent. We compute the null hypothesis by jittering the exact timing of each spike within spike triplets. Subtraction of the null hypothesis from the measured synchrony provides a rate-corrected measure of synchrony, equivalent to the covariance between the two neurons, as a function of time during pursuit. Deviations of the covariance curve from zero would indicate changes in synchrony that are unexpected given the instantaneous firing rates of the two neurons under study.

### Statistical analysis

We used the Julia package HypothesisTests for common statistical calculations, including t-tests for correlation and independent samples t-tests. All statistical tests were two-sided and we report exact p-values where possible. To perform permutation tests, we randomly sampled with replacement from two comparison populations under the null hypothesis. We performed 10,000 random permutations for each test unless otherwise noted.

## Data availability

All data analyzed for this study have been deposited into the Open Science Framework database (doi: https://doi.org/10.17605/osf.io/xwmah). Code for the models is available via Github (https://doi.org/10.5281/zenodo.19685373). Requests for additional data or analyses can be made to the corresponding author.

## Acknowledgements

We thank Stefanie Tokiyama and Bonnie Bowell for assistance with animal care. We thank Court Hull and Nuo Li for helpful comments on an earlier version of the manuscript. We also thank Nathan J. Hall and Seth W. Egger for helpful comments and discussions.

## Author contributions

DJH and SGL designed all experimental and analysis procedures. DJH performed extracellular recordings in the cerebellar circuit. DJH analyzed the data. DJH and SGL generated the figures and wrote the manuscript.

