## Extended Data Figures 1-8 for "Circuit mechanisms to transform neural population dynamics for motor control"

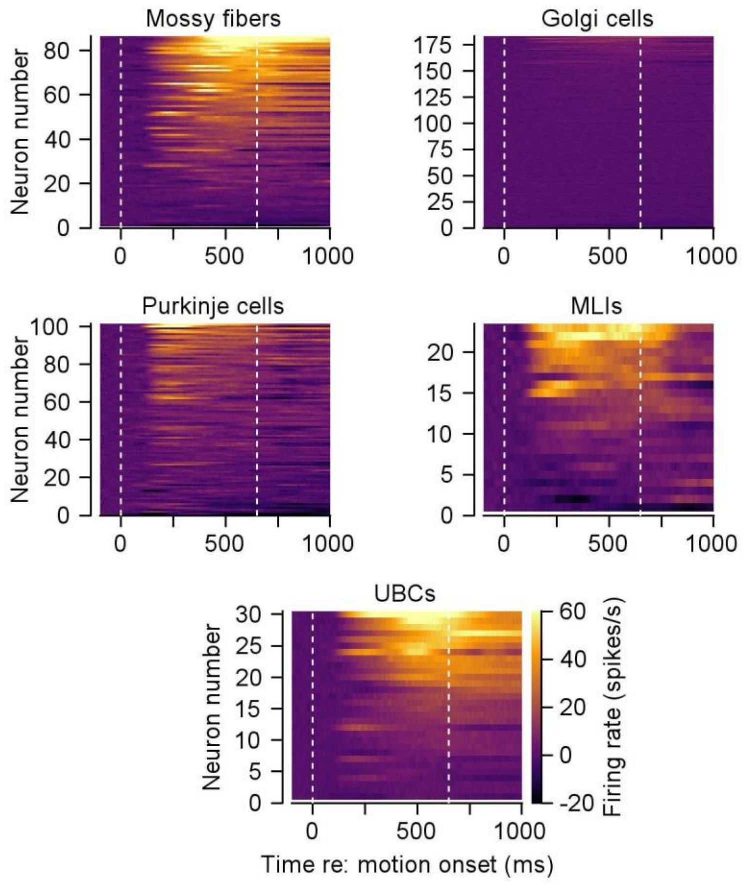


**Extended Data Fig. 1. Summary of the responses of all expert-identified floccular neurons during pursuit.** Each heatmap plots the responses of each neuron in our sample in its preferred direction as a function of time during pursuit. Each row in the heatmap corresponds to a single neuron. The two vertical dashed lines in each heatmap indicate the time of target motion onset and offset. The color axis is shared across all heatmaps. Note several features of the responses: 1) in all neuron types, there is a range of responsiveness with some neurons of each type responding weakly or not at all; 2) Golgi cells are largely unresponsive; 3) mossy fibers and unipolar brush cells (UBCs) that respond during pursuit tend to remain active at the end of the trial, after target motion has ceased; 4) Purkinje cells and molecular layer interneurons (MLIs) that respond during pursuit tend to return to baseline firing at the end of the trial, after target motion has ceased. In evaluating the population responses, it is important to understand our sampling procedure. Unlike recordings made with single microelectrodes in previous studies, where we adjusted the depth of the electrode until we had isolated the spikes from a single highly-responsive neuron, in this study we simply positioned a multi-contact probe at a location where there was eye movement activity. We then recorded from all neurons on all contacts, sorted spikes, and included all isolated neurons in our sample whether they responded strongly, weakly, or not at all.

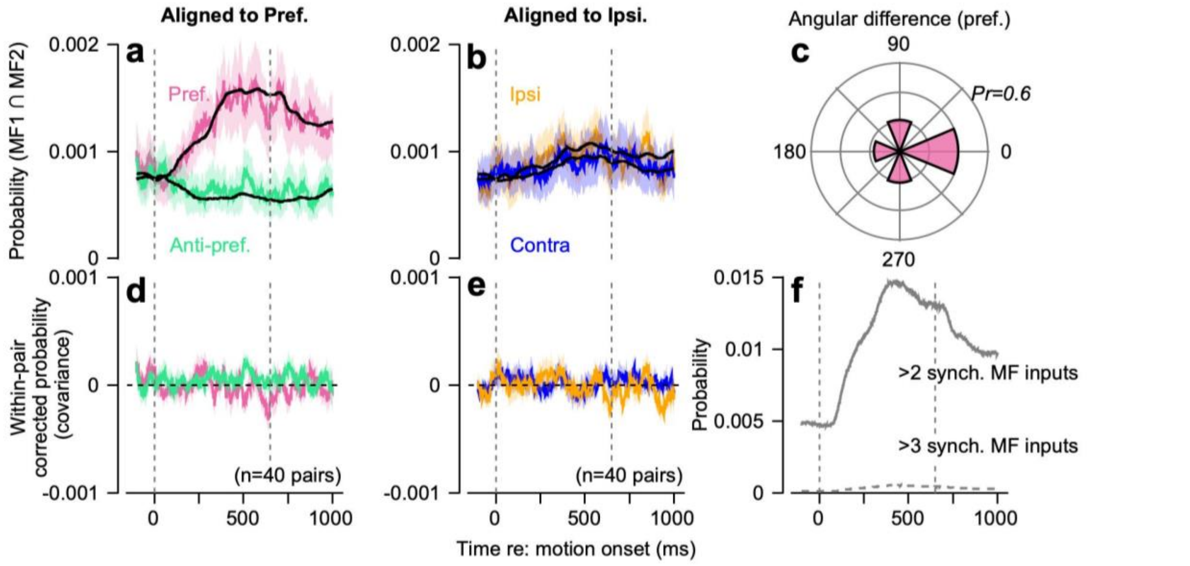


**Extended Data Fig. 2. Evaluation of millisecond scale synchrony of simultaneously-recorded mossy fibers. a,** Mean probability of two simultaneously-recorded mossy fibers firing together in the same millisecond, across all recorded simultaneously recorded pairs. Colored curves show probability in the preferred and anti-preferred directions of one mossy fiber, where the direction chosen for alignment was randomly selected from the pair. Black lines denote the probability of synchronous spikes that would be expected given the firing rates of two statistically independent neurons. **b,** Same as in **a**, but aligned to the ipsiversive pursuit direction. **c,** Angular difference between the preferred directions of the pairs of simultaneously-recorded mossy fibers. **d,** Within-pair corrected probability for the two-colored traces shown in **a** (colored curve minus black curve). **e,** Same as in **d**, but aligned to the ipsiversive pursuit direction. **f,** Mean probability of observing millisecond-scale synchrony between four randomly-selected mossy fibers from the complete population, aligned to their preferred directions. Solid and dashed gray curves show probability of 2 or more, or 3 or more, synchronous spikes in a given millisecond. As mossy fiber pairs show no millisecond scale synchrony, the probability of multiple mossy fibers firing together follows the same profile as the mean mossy fiber response.

3

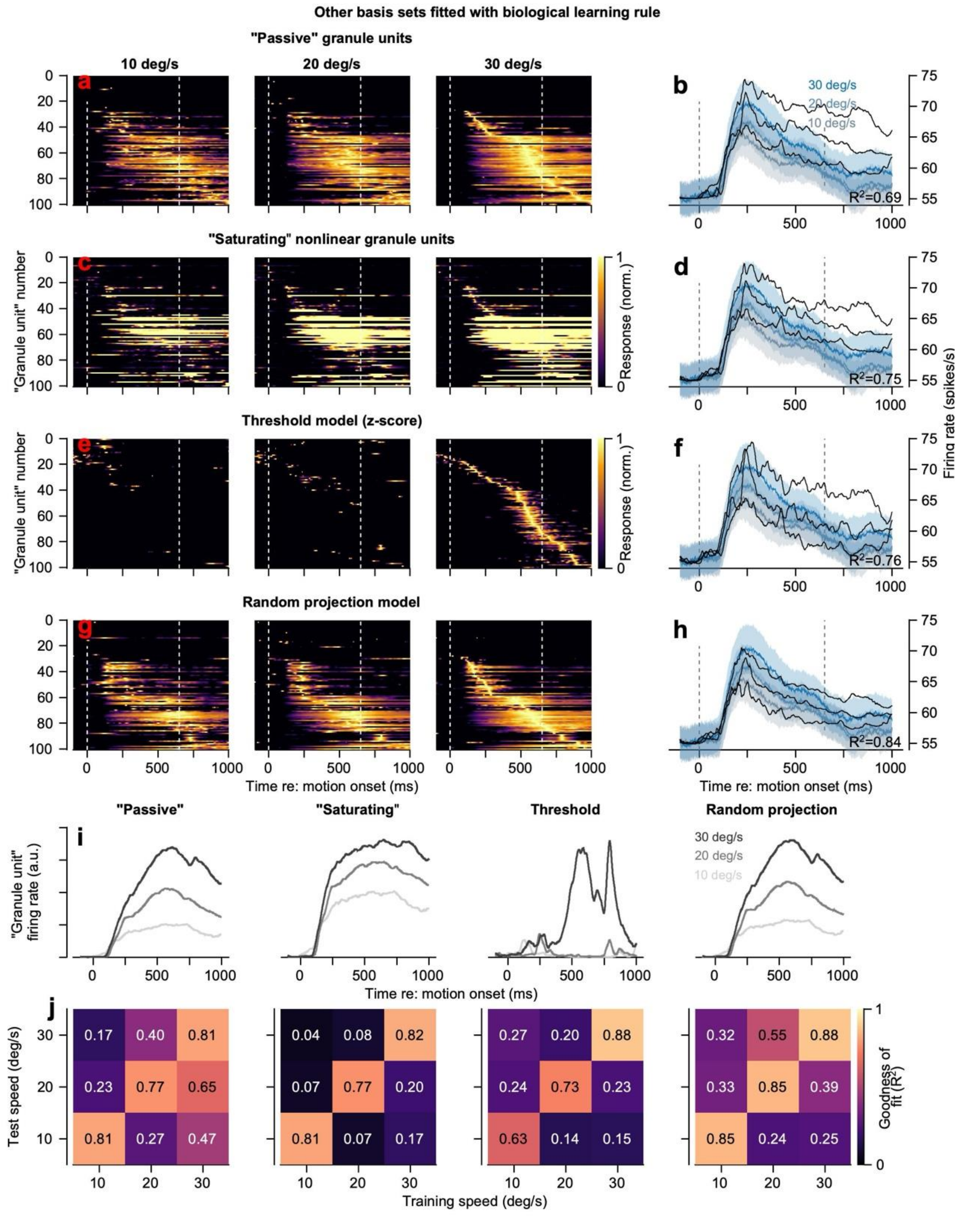


**Extended Data Fig. 3. Characteristics of alternative granule cell basis set responses to mossy fiber inputs using the LTP/LTD learning rule for parallel fiber to Purkinje cell weights. a, c, e, g,** Responses of "granule units" in models that contained no dynamics (**a**), a compressive non-linearity (**c**), an input-dependent threshold[90] (**e**), or recurrent inhibition from random connections[41,57,91,92] (**g**). The three heat maps in each row show responses to mossy fiber inputs across three pursuit speeds (10, 20, and 30 deg/s). Each granule unit's response was normalized to its peak response across all three pursuit speeds and ordered based on the timing of the peak response to 30 deg/s target motion. **b, d, f, h,** Best fit to Purkinje cell activity in the preferred direction for each granule unit population response. Different shades of blue show responses for different pursuit speeds. The Pearson $R^2$ value is shown for each fit, averaged across all neurons. **i,** Mean granule unit population responses for each basis set model across pursuit speeds. **j,** Generalization matrices showing the $R^2$ for each model when trained to predict Purkinje cell responses from a single pursuit speed and tested on responses from all other speeds.

All four models of granule unit activity form reasonable basis sets to support Purkinje cell firing that has roughly the correct shape and scales with pursuit speed. However, inspection of the model fits for the "Passive", "Saturated" and "Thresholded" models shows that the optimized models could not capture the time course of Purkinje cell firing very well, even though the $R^2$ values exceeded 0.83. The random projection model allows excellent fits to the time course of Purkinje cell firing, but it requires active computation by inhibitory elements within the granule layer (e.g., Golgi cells) that inactivate granule unit responses across time. Our data shows limited evidence for Golgi cell involvement in granule representations, leading us to discount this model of granule layer activity at least in the floccular complex. Each of the granule unit models generalizes fairly well to other speeds when trained on a single speed, but none as well as the STP granule unit population.

The most important observation from the granule unit basis sets is that optimization allowed the model to set thresholds on granule unit activity in a way that made their responses transient and allowed them to distribute across the duration of a behavioral trial. Then, the optimization strategy was able to weight the granule unit activities to yield reasonable fits to the Purkinje cell firing. Even though all the basis sets are derived by thresholding granule units in a way that treads on the noise in responses, they each contrived to create qualitatively similar granule unit basis sets. We conclude that the temporal transformation performed in the floccular complex is performed most easily by temporal decomposition of mossy fiber inputs. Cleaner decompositions of time lead to better performance. Because the STP mechanism described in the main figures and text is biologically plausible, well supported by cellular analyses, and treads on the signal in the mossy fiber inputs rather than the noise, we prefer to think that it is a better representation of reality. In the final analysis, however, the main lesson from the analysis here is that the properties of the granule unit basis set are critical for the success of the computation.

4

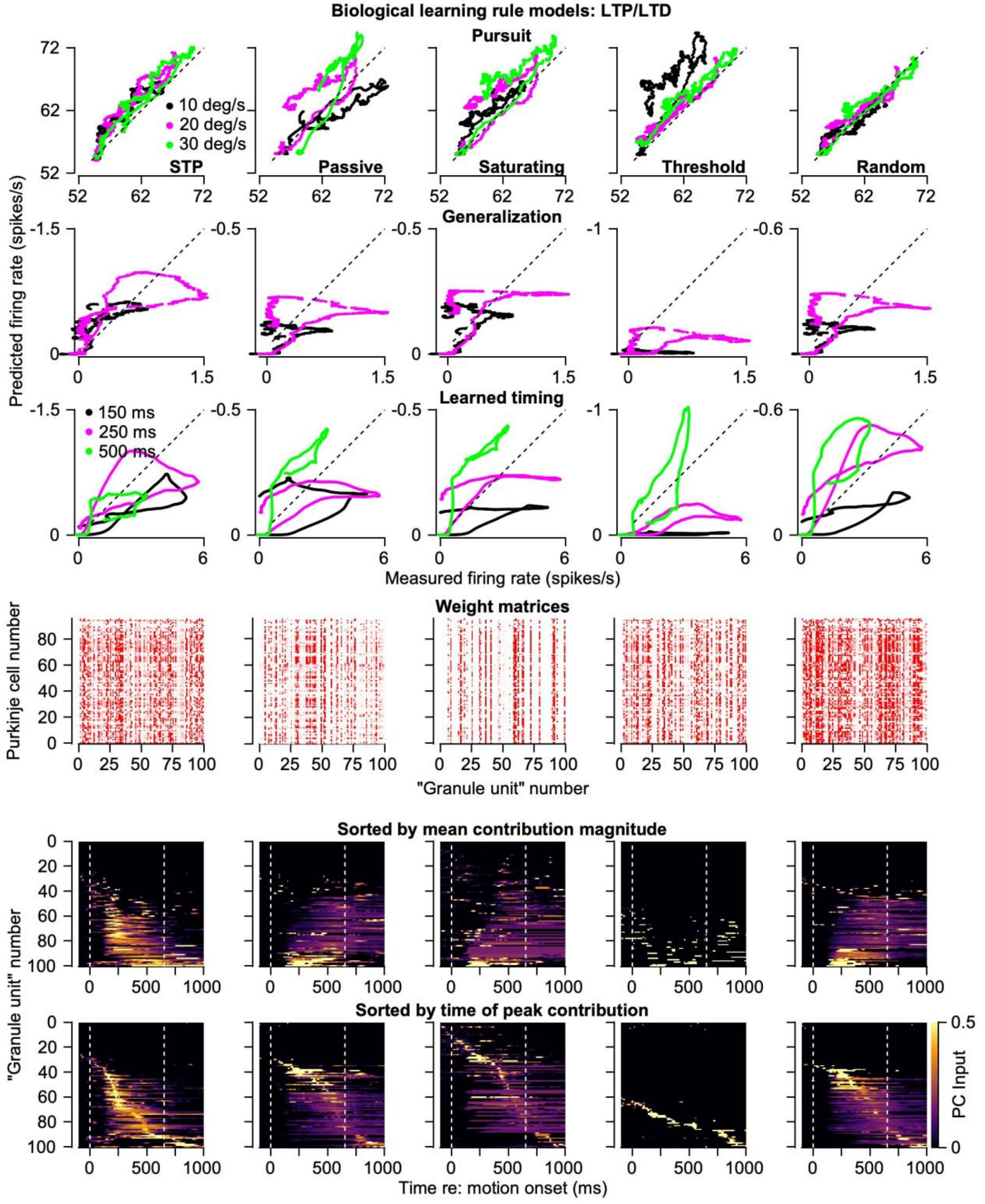


**Extended Data Fig. 4. Quantitative analysis of performance of alternative granule cell basis sets using the LTP/LTD learning rule for parallel fiber to Purkinje cell weights.** Each column of plots shows data for a different granule unit basis set. From left to right: STP, linear, saturating non-linearity, threshold, and random projection. From top to bottom, the first three rows plot the predicted firing rate from the optimized model as a function of the measured firing rate, where each symbol represents a different time starting 100 ms before the onset of target motion. The first row shows data for models optimized only to pursuit in the preferred direction, different colors show different target speeds; second and third rows show data for the generalization of pursuit learning to different target speed and temporal specificity of learning for models optimized for pursuit in all directions and including both molecular layer interneuron and granule unit inputs to Purkinje cells. Note that the magnitude of modeled learned responses in the second and third rows are arbitrary, and the graphs have been scaled to be subjectively similar. Fourth row shows the matrices of weights from granule units to Purkinje cells for the preferred direction models, normalized so that the colors run from zero to the value that would reflect a uniform distribution of weights across 100 granule units (0.01). In the 5 models, from left to right, 67.3%, 64.1%, 78.2%, 70.6%, and 64.8% of the 9600 weights were non-zero. Heatmaps in the fifth and sixth rows show the mean contributions to the full population of Purkinje cell firings, where each horizontal line is a different granule unit as a function of time. Granule units are ordered according to the magnitude of the input to Purkinje cells in the fifth row and according to the time of the peak input in the sixth row. In the sixth row, granule units were placed at the top of the heatmap if the z-score of their contribution to Purkinje cell firing was less than 0.1.

Subjectively, the STP model performs best across the first 3 rows, but all the models perform somewhat acceptably. Ideal performance in all 3 rows would be represented by all points plotting along the diagonal dashed line. For reconstruction of Purkinje cell firing across pursuit speeds (1st row) only the STP and random projection models show excellent performance across all three tested pursuit speeds. For generalization to target speed (2nd row), the STP model comes closest to plotting along the diagonal although all models fail to some degree for 20 deg/s pursuit speeds. For learned timing (3rd row), the STP model shows concentric responses that come close to the diagonal while the other models reproduce learned timing less well. The weight matrices (4th row) tend to have a vertical appearance, meaning that some granule units were used little or not at all. The heatmaps in the 5th and 6th rows reveal how each model worked and why they all were able to fit at least the firing during pursuit fairly well. The STP model clearly used more of the granule units than the other models (5th row) while the Threshold model used very few. Inspection of the 6th row shows that all of the models optimized parameters to produce granule unit basis sets that had transient responses distributed across the time of the trial. In particular, they optimized firing thresholds in a way that treads on noise in the input mossy fibers and thus created a basis set that performed the temporal input-output transformation in our data. All except the STP basis set failed to some degree on generalization and learned timing because the optimization strategy used different granule units for model responses to different speeds, and because the coverage of the second half of the trial was weak. We take the reasons for success and failure of the non-STP basis sets as evidence that the appropriate basis set, no matter the mechanism that creates it, has two critical features: (1) temporal decomposition of sustained inputs in a way that tiles the during of a trial and (2) scaling and consistent timing of each granule unit's responses across target speeds.

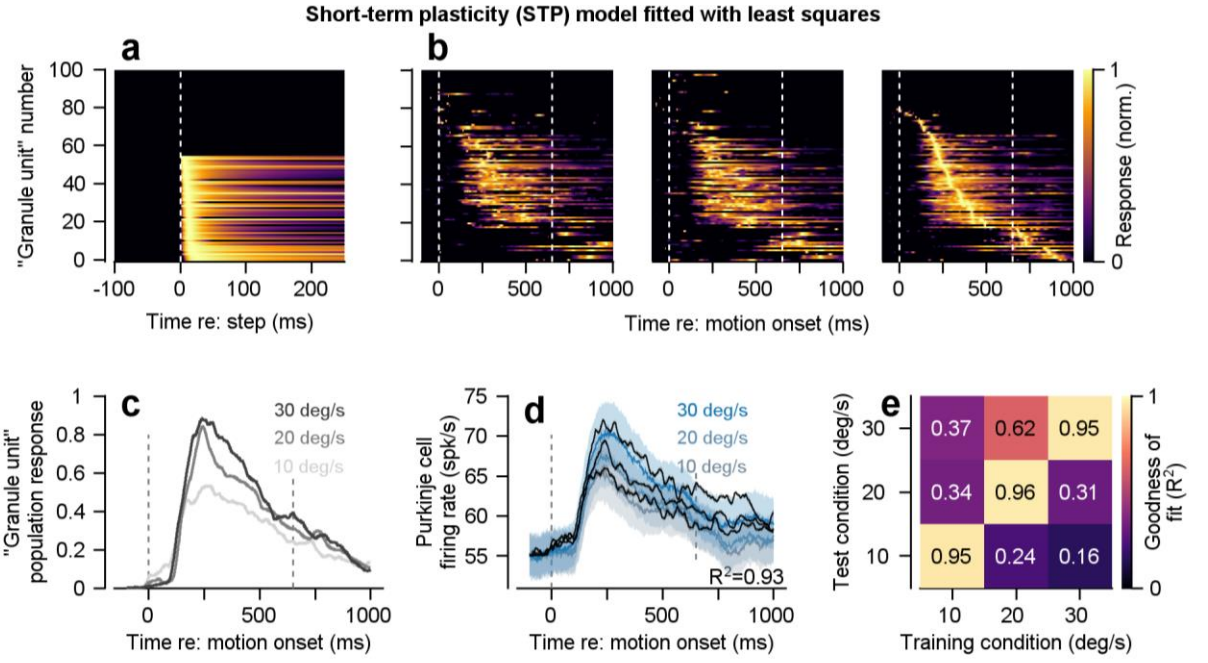


**Extended Data Fig. 5. Performance of the short-term plasticity (STP) granule unit basis set using least squares optimization of parallel fiber to Purkinje cell weights for preferred-direction responses.** The panels are meant to parallel the bottom half of Fig. 3, using the same STP strategy for the granule unit basis set but optimizing the weights of parallel fiber to Purkinje cell synapses using non-negative least squares instead of the LTP/LTD learning rule. **a,** Responses of granule units to a step input of mossy fiber responses in a model based on short-term plasticity (STP) in the mossy fiber to granule unit synapses. **b,** Responses of granule units in the STP model to four randomly sampled mossy fiber inputs across pursuit speeds. **c,** Population responses of granule units in the STP model. **d,** Fits (black traces) to the mean responses of Purkinje cells across pursuit speeds (colored traces) using least squares to adjust granule unit weights on Purkinje cells. **e,** Generalization matrix showing the model $R^2$ when trained on the data for one target speed and tested on all 3 target speeds. Note that generalization compared to Fig. 3i is significantly worse, owing to the concentration of non-zero parallel fiber-to-Purkinje cell weights on a small subset of granule units, described further in Extended Data Fig 7.

6

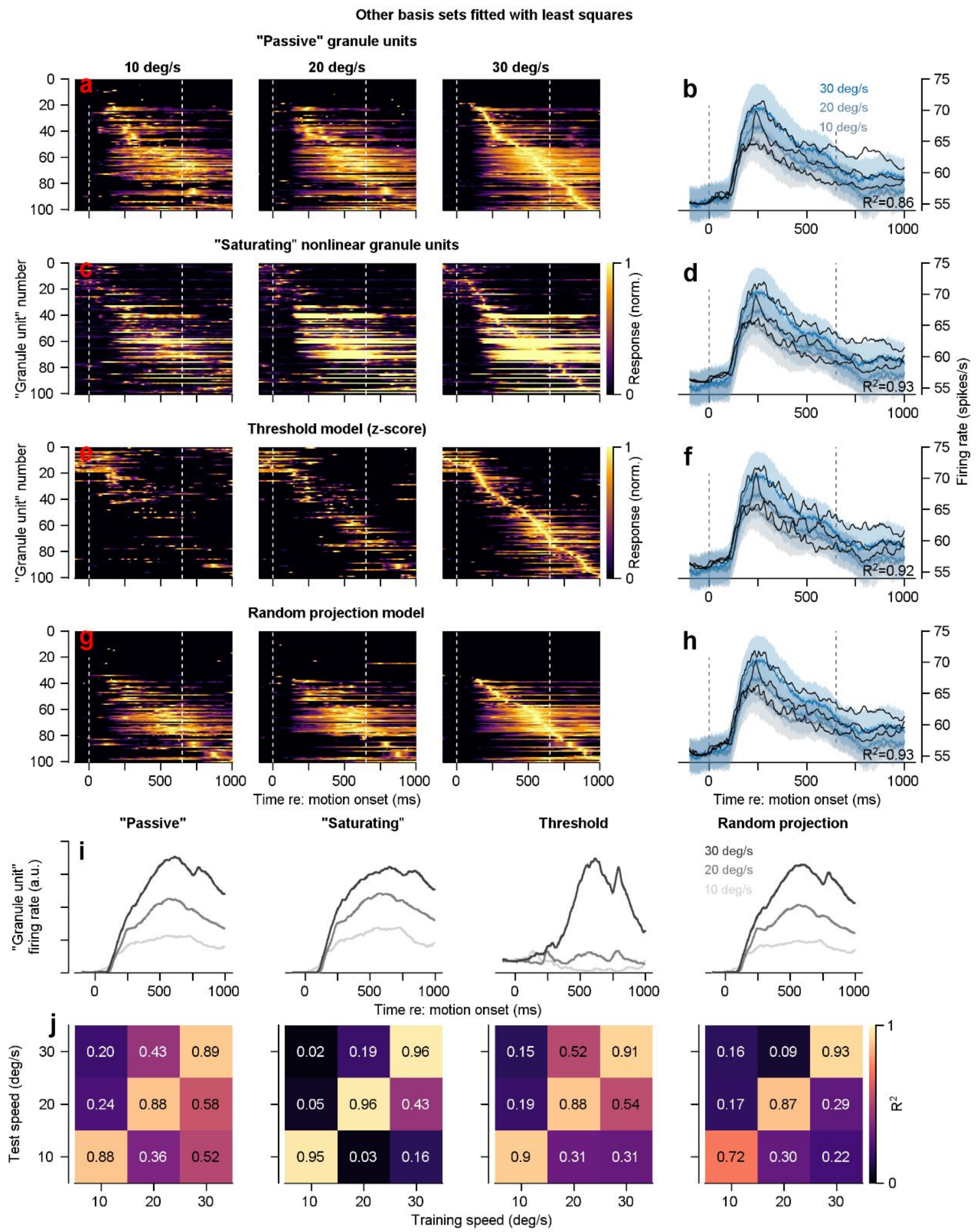


**Extended Data Fig. 6. Characteristics of alternative granule cell basis set responses to mossy fiber inputs using least squares to adjust parallel fiber to Purkinje cell weights for preferred-direction responses.** Panels in this figure are identical to Extended Data Fig. 3, with only difference being how the parallel fiber-to-Purkinje cell weights are determined. In Extended Data Fig. 3, a biological plausible LTP/LTD learning rule was used. In the current figure, we use a non-negative least squares approach. The most important observation is that all models perform well using a least squares rule, which is enabled by selecting a relatively small subset of transient granule units tiled across the trial. While reproduction of Purkinje cell responses is excellent for all models (**b, d, f, h**), selection of a diminutive set of granule units results in impaired generalization performance (**j**) in all models compared to the LTP/LTD learning rule.

7

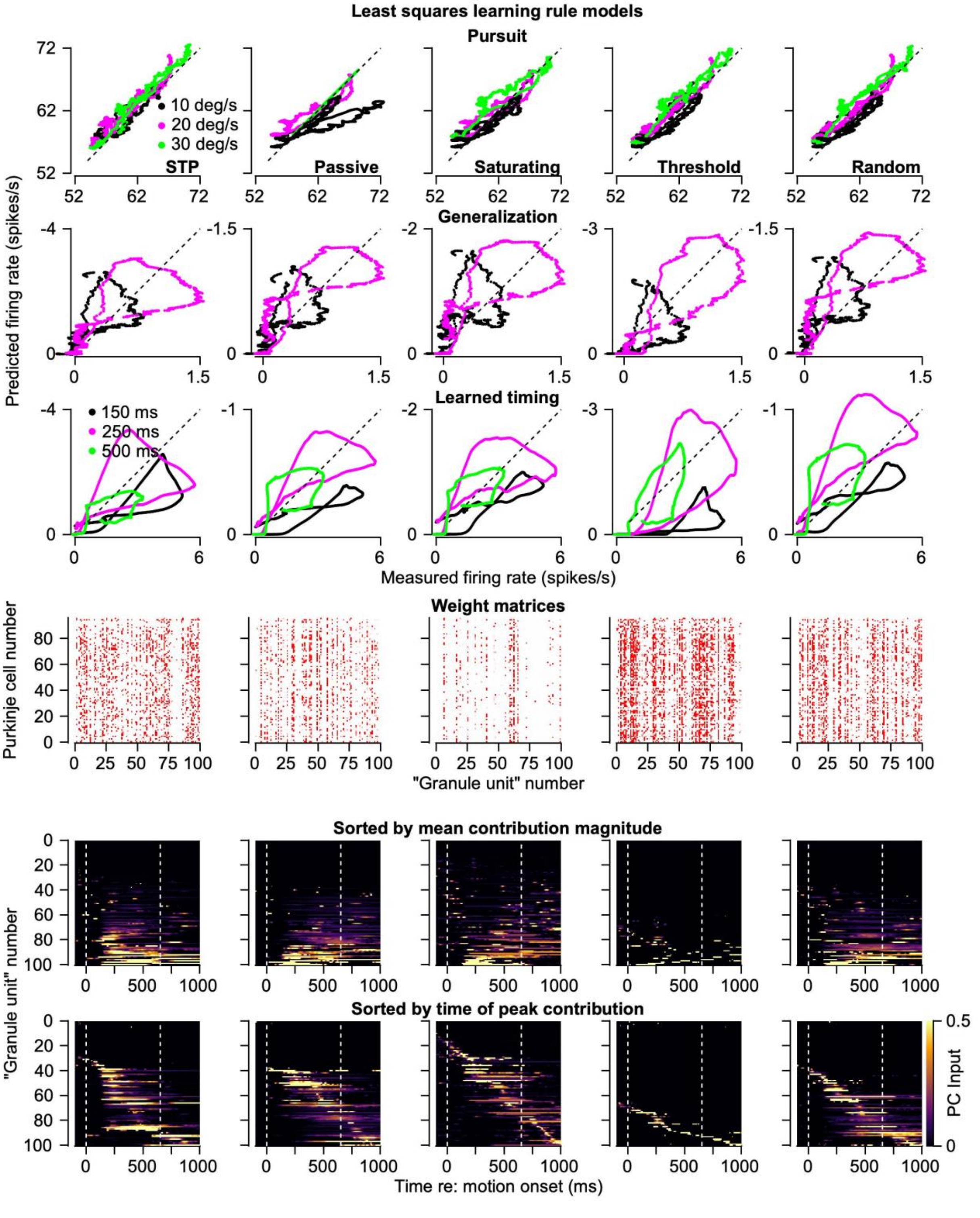


**Extended Data Fig. 7. Quantitative analysis of performance of alternative granule cell basis set using least squares to adjust parallel fiber to Purkinje cell weights.** Each column of plots shows data for a different granule unit basis set. From left to right: STP, linear, saturating non-linearity, threshold, and random projection. From top to bottom, the first three rows plot the predicted firing rate from the optimized model as a function of the measured firing rate, where each symbol represents a different time starting 100 ms before the onset of target motion. From top to bottom: first row shows data for models optimized only to pursuit in the preferred direction, different colors show different target speeds; second and third rows show data for the generalization of pursuit learning to different target speed and temporal specificity of learning for models optimized for pursuit in all directions and including both molecular layer interneuron and granule unit inputs to Purkinje cells. Note that the magnitude of modeled learned responses in the second and third rows are arbitrary, and the graphs have been scaled to be subjectively similar. Fourth row shows the matrices of weights from granule units to Purkinje cells for the preferred direction models, normalized so that the colors run from zero to the value that would reflect a uniform distribution of weights, 0.01. In the 5 models, from left to right, 18.4%, 14.4%, 19.3%, 29.2%, and 20.6% of the 9600 weights were non-zero. Heatmaps in the fifth and sixth rows show the mean contributions to the full population of Purkinje cell firings, where each horizontal line is a different granule unit as a function of time. Granule units are ordered according to the magnitude of the input to Purkinje cells in the fifth row and according to the time of the peak input in the sixth row. In the sixth row, granule units were placed at the top of the heatmap if the z-score of their contribution to Purkinje cell firing was less than 0.1.

Subjectively, the STP model performs best across the first 3 rows, but all the models perform somewhat acceptably. Ideal performance in all 3 rows would be represented by all points plotting along the diagonal dashed line. For reconstruction of Purkinje cell firing across pursuit speeds (1st row), except for the passive model, all models show excellent performance across all three tested pursuit speeds. For generalization to target speed (2nd row), the STP model comes closest to plotting along the diagonal although all models fail to some degree for 20 deg/s pursuit speeds. For learned timing (3rd row), the STP model shows concentric responses that come close to the diagonal while the other models reproduce learned timing less well. The weight matrices (4th row) tend to have a vertical appearance, meaning that some granule units were used little or not at all. The heatmaps in the 5th and 6th rows reveal how each model worked and why they all were able to fit at least the firing during pursuit fairly well. The STP model clearly used more of the granule units than the other models (5th row) while the Threshold model used very few. Inspection of the 6th row shows that all of the models optimized parameters to produce granule unit basis sets that had transient responses distributed across the time of the trial. In particular, they optimized firing thresholds in a way that treads on noise in the input mossy fibers and thus created a basis set that performed the temporal input-output transformation in our data. All except the STP basis set failed to some degree on generalization and learned timing because the optimization strategy used different granule units for model responses to different speeds, and because the coverage of the second half of the trial was weak. We take the reasons for success and failure of the non-STP basis sets as evidence that the appropriate basis set, no matter the mechanism that creates it, has two critical features: (1) temporal decomposition of sustained inputs in a way that tiles the during of a trial and (2) scaling and consistent timing of each granule unit's responses across target speeds.

Comparison with Extended Data Figure 4 reveals that the least squares learning rule used many fewer granule units than did the LTP/LTD learning rule. The weight matrices in the 4th row and the heatmaps in the 5th and 6th rows reveal cherry-picking of the parallel fibers that best reproduce the required temporally decomposed basis set, which we consider to be non-biological.

8

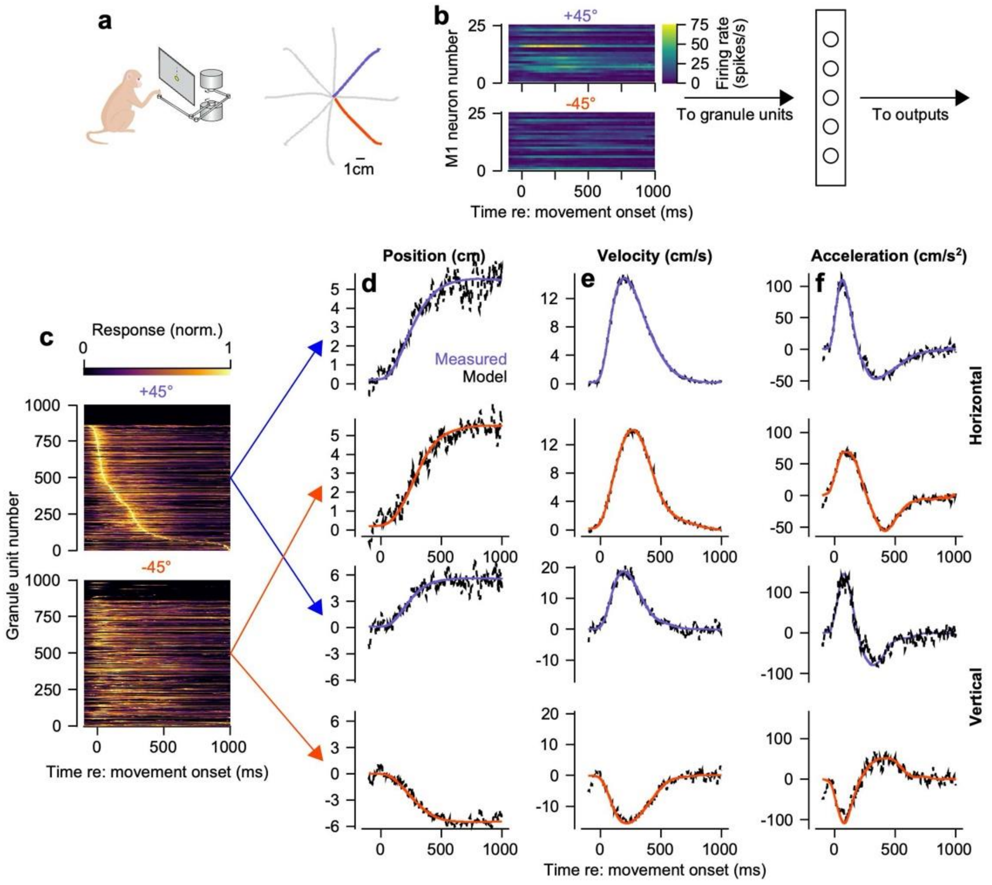


**Extended Data Fig. 8. Granule basis set responses enable transformation of primary motor cortex population dynamics into kinematic outputs during reaching movements. a,** Schematic diagram of the reaching task (left) and reach trajectories to eight peripheral targets (right), with the +45 and –45 degree directions highlighted in blue and red. **b,** Heatmap of motor cortex unit activity during reaching movements[47,48]; for visualization, 25 units were randomly selected from the population of $n = 573$ neurons. **c,** Heatmaps show the activity of 1,000 simulated granule cell units as a function of time during center-out reaching in the two highlighted directions. **d-f,** Decoding of limb kinematics (position, velocity, and acceleration) in both directions as a function of time. Reconstructed kinematic signals are shown as dotted black lines. Panel organization matches that of Fig. 6.

9
